# Cryo-electron tomography suggests tubulin chaperones form a subset of microtubule lumenal particles with a role in maintaining neuronal microtubules

**DOI:** 10.1101/2022.07.28.501854

**Authors:** Saikat Chakraborty, Antonio Martinez-Sanchez, Florian Beck, Mauricio Toro-Nahuelpan, In-Young Hwang, Kyung-Min Noh, Wolfgang Baumeister, Julia Mahamid

## Abstract

The functional architecture of the long-lived neuronal microtubule (MT) cytoskeleton is maintained by various MT-associated proteins (MAPs), most of which are known to bind to the MT outer surface. However, electron microscopy (EM) has long ago revealed the presence of particles inside the lumens of neuronal MTs, of yet unknown identity and function. Here, we use cryogenic electron tomography (cryo-ET) to analyze the three-dimensional (3D) structures and organizations of MT lumenal particles in primary hippocampal neurons, human induced pluripotent stem cell-derived neurons and pluripotent P19 cells. We obtain in-cell 3D maps of several lumenal particles from the respective cells and detect structural features that are common to all cell-types, underscoring their potential overarching functions. Mass spectrometry-based proteomics combined with structural modeling suggests a subset of lumenal particles could be tubulin-binding cofactors (TBCs) bound to tubulin monomers. A different subset of smaller particles, which remains unidentified, exhibits densities that bridge across the MT protofilaments. We show that increased lumenal particle concentration within MTs is concomitant with neuronal differentiation and correlates with higher MT curvatures. Enrichment of lumenal particles around MT lattice defects and at freshly polymerized MT open-ends suggest a MT protective role. Together with the identified structural resemblance of a subset of particles to TBCs, these results hint at a role in local tubulin proteostasis for the maintenance of long-lived neuronal MTs.

## Introduction

The microtubule (MT) cytoskeleton plays an essential role in neuronal morphogenesis (1), supporting axonal trafficking (2), signal transduction (3), axon guidance (4) and synapse formation (5). Irregularities in MTs lead to abnormal morphogenesis and ultimately to detrimental neurodevelopmental defects (6–8). Thus, the neuronal MT cytoskeleton is maintained by complex layers of regulatory mechanisms that include a pool of neuron-specific MT-associated proteins (MAPs) (9). These MAPs predominantly bind to the outer surface of the hollow MT lattice and regulate the dynamics as well as material properties of the MT cytoskeleton (10–12). Early conventional EM studies further showed the presence of periodically arranged particles within the MT lumen of insect epithelia (13), spermatids (14) and blood platelets (15). Cryo-electron tomography (cryo-ET) studies have since unambiguously confirmed the presence of such lumenal particles in unstained, frozen-hydrated sections of different cells, and showed their high abundance particularly in neurons (16–25). However, lumenal particles were not detected inside MTs polymerized *in vitro* from purified brain tubulins (17). These studies suggest that the *in vivo* repertoire of neuronal MT-associated proteins could be more complex than perceived; to date our knowledge about the structures, molecular identities or functions of neuronal lumenal particles remains limited.

The MT lumen, with a diameter of approximately 17 nm, is largely excluded from the cytoplasm except for the two filament ends and lateral openings that form upon regulated severing and/or defects (26, 27). Theoretical modeling predicts very slow diffusion of molecules with binding affinity for tubulins through the MT lumen (28). Nevertheless, regulation of MT stability by selective localization of MAPs or posttranslational modification (PTM) in the MT lumen may provide a favorable mechanism, since these processes should not interfere with the multitude of trafficking events happening on the MT outer surface (29). Such a mechanism is indeed supported by findings of MT inner-proteins (MIPs) in extraordinary stable cortical MTs in parasites and specialized doublet MTs in sperm tails and cellular appendages such as cilia and flagella that maintain the structural integrity of the MT lattice during extreme mechanical deformations caused during propulsive motion (30–34). However, the roles of MIPs in cytoplasmic MTs remain poorly described. Exceptions are studies showing acetylation of lumenal K-40 of alpha-tubulin by Tubulin Acetyltransferase (TAT) that increases MT mechanical resilience, lattice integrity and longevity (35–37). Not surprisingly, acetylated MTs are an integral part of axonal MT bundles that function as highways for intracellular transport (38). Therefore, TATs have been suggested to be a component of the lumenal particles (39), as well as deacetylases such as HDAC6 that access the MT lumen by hitchhiking on the MT plus-end tracking protein EB1 for removing lumenal acetylation marks (40, 41). A recent report suggests that MAP6 could also reside in the MT lumen, and perform similar functions as MIPs in MT doublets by stabilizing a coiled architecture of neuronal MTs (42). However, there is no direct structural evidence to show that these proteins are components of MT lumenal particles in neurons. Thus, a major question remains about the identities and functions that such abundant particles preform in the neuronal microtubule cytoskeleton.

In this study, we utilized *in situ* cryo-ET to elucidate the molecular architectures and nanometer-scale organization of lumenal particles in three vitrified neuronal cell types: rodent primary hippocampal neurons, human induced pluripotent stem cell (hiPSC)-derived neurons, and murine pluripotent P19 cells – a neuronal precursor cell line used as model system for neuronal differentiation. We developed an approach comprising automated particle detection, subtomogram averaging (STA), and spatial statistics combined with mass spectrometry-based proteomics to interrogate structures and potential functions of the lumenal particles in their native context. Our data provide important clues as to the particles identity, show that MT lumenal particles are an integral part of the differentiated neuronal MT cytoskeleton, and that they may be involved in MT quality control.

## Results

### Organization of lumenal particles within MTs

Cryo-ET of vitrified intact thin cellular processes (thickness <0.2 μm) of rodent primary hippocampal neurons, murine pluripotent P19 cells and hiPSC-derived neurons revealed MT bundles (**Fig. 1A-F, SI Appendix, videos S1-S3**). Globular particles were densely packed within the MT lumens of the primary (17) and hiPSC-derived neurons (**Fig. 1D-G**). In comparison, particles were more sparsely distributed in the murine pluripotent P19 cell line (**Fig. 1G**). Close inspection showed morphological similarity among lumenal particles across the cell types (arrowheads; **Fig. 1G**). Among those, the most notable feature was the presence of a ring-shaped particle, which was also recently described inside mouse dorsal root ganglion (DRG) and hippocampal neuronal MTs (22, 23).

**Figure 1:**
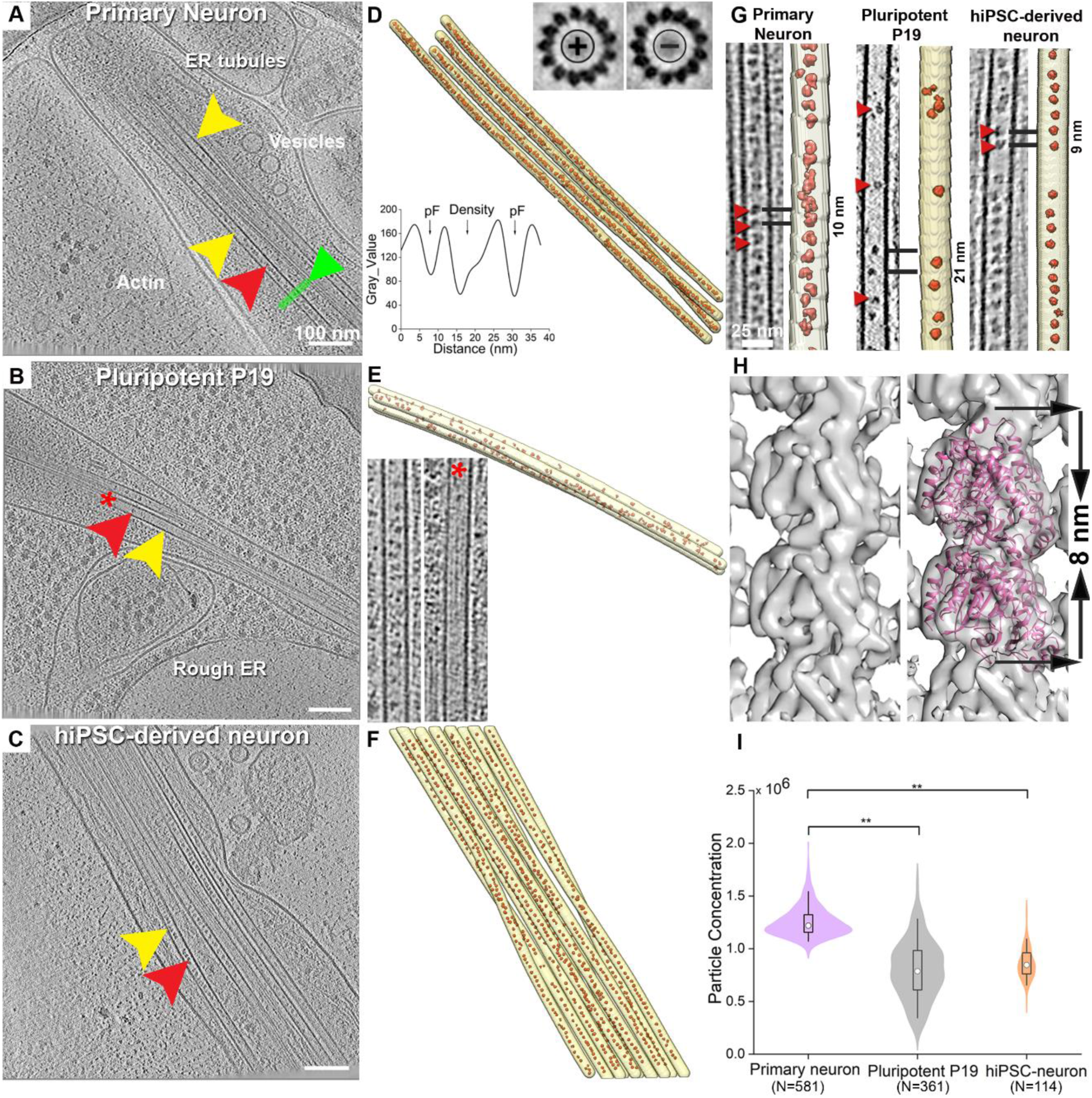
Organization of lumenal particles inside neuronal MTs. **A-F**. Representative tomographic slices and 3D-surface rendering of traced MTs (yellow arrowhead), along with the detected lumenal particles (red arrowhead) of indicated cell types: **A**, **D**. Rodent hippocampal primary neuron, slice thickness 9 nm; Insets in D: Top right, subtomogram averages of individual MTs shows different polarity. Below left: line profile across a MT indicated by green arrowhead in A shows presence of extra density between pFs; **B**, **E**. Pluripotent P19 cell, slice thickness 6.8 nm; Inset: Zoomed in image of an empty MT marked by red star in (shown at different orientation) compared to its neighboring MT. **C**, **F**. hiPSC-derived neuronal process, slice thickness 4.25 nm. **G**. Tomographic slices showing varying particle abundances in the indicated cell types. Segmentations of the MTs (Yellow) and lumenal particles (red) are shown. Typical distances are indicated. **H.** 8.2 Å *in situ* subtomogram average of the 13 pF rodent primary neuronal MTs, and with fitted tubulin dimer (right). Dimension of a tubulin dimer indicated. **I.** Quantification of the particle concentrations (numbers/µm^3^ of lumenal volume) in indicated cell types represented as boxplot within a violin. Median values are marked by the white circle. Asterisks indicate Mann-Whitney test significance: ** p<0.01. N, number of MTs analyzed. See also Figure S1.

We established the MT polarities in the imaged cellular processes of primary neurons using STA (43). STA provided a 8.2 Å *in situ* map of primary neuronal MTs (**Fig. 1H, S1A-I**) with discernible secondary structural features showing that MTs exclusively constituted of 13 pFs (**Fig.1H, S1H**) and conforming to the consensus lattice architecture observed in mammals (44). We found both mixed (Inset: **Fig. 1D, Fig. S1A-C, SI Appendix)** and uniform polarity orientations (**Fig. S1D-F)** in different processes, which indicate their identities as dendrites or axons, respectively (45). Lumenal particles were present within all MTs irrespective of their polarity **(Fig. S1B, E)**.

To quantify the abundances of the lumenal particles, we developed an automated method for template-free detection of particles inside the segmented MT lumens (**SI Appendix, Fig. S2A-C**, see methods) by employing discrete Morse theory segmentation and topological persistence simplification (46). In accordance with the visual inspection, the analysis revealed the highest concentration of lumenal particles in the primary neurons, as noted in earlier studies (17, 24), followed by that in hiPSC-derived neurons (**Fig**. **1I**). In case of pluripotent P19 cells, particle concentration distribution was broad, with neighboring MTs containing markedly different concentrations (inset: **Fig. 1E**).

Next, we employed nearest-neighbor (NN) analysis (46) to describe the organization of the lumenal particles using the particle coordinates refined in RELION (47) (described below). We observed non-random distributions in all samples, with peaks at 8-10 nm representing most common NN-distance (black dashed line, **Fig. S2D-F**) (17, 21, 25). This suggested significant short-range order of particles (≤10 nm), with the highest probability observed in primary neurons. Longer distances were not statistically significant, as they were also obtained from simulations of randomly distributed particles (gray shaded area, **Fig. S2D-F**). We statistically evaluated these spatial patterns for higher order organization at multiple distance scales using Ripley’s L function against complete randomness with numerical corrections for cryo-tomograms (46)(**Fig. S2G**-**I**, **SI Appendix**). Here, positive L values indicate clustering, and negative values indicate uniform distribution (given it is separated from complete randomness). We observed negative L values for lumenal particles, signifying their uniform distribution in the primary and hiPSC-derived neurons. However, L values for P19 cells overlap within the complete randomness indicating that here lumenal particles are randomly distributed. Our data and analyses thus agree with previously reported organizational differences of lumenal particles observed across different cell types (16, 17, 21, 22).

### 3D morphologies of lumenal particles

Next, we sought to elucidate the structures of the lumenal particles for each cell type using STA (**Table S1**). Marked structural and compositional heterogeneity, combined with the relatively small size of the particles (diameters in the range of 7-10 nm), posed a challenge for STA-based structure determination. In order to generate homogeneous particle groups for averaging, we subjected the particles to several sequential rounds of 3D classifications until no new classes emerged (**SI Appendix**, **Fig. S3)**. All class averages were obtained *de novo*, *i.e.* by reference-free analysis, validated against different references, and contained sufficient angular sampling (**Fig. S4A-G**). The approximate molecular masses of the lumenal particles were derived from the tomograms using the ribosomes as an internal reference, and found to be in the range of 200-400 kDa (**SI Appendix, Fig. S5A-F**), in agreement with a previous analysis (22). The size and molecular mass of these particles safely precluded a classical motor–cargo complex that can have molecular mass up to several mega Daltons (48).

Visual analyses of the pluripotent P19 tomograms suggested two types of particles, based on their association with the MT wall; a subset of the particles showed connections with the MT wall, while others appeared floating inside the lumen (**Fig. 2A**, tomogram slice). STA supported the visual inspection (**Fig. 2A**). In the MT-wall bound category, the most notable class averages were elongated particles with four domains arranged in a crisscross fashion. We termed this density - ‘bound MT inner protein 1’ or bMIP1, and bMIP2, which we found to attach to the MT wall with stalk-like densities (pink arrowheads, **Fig. 2A**). The distance between the neighboring stalks was ∼4 nm, equivalent to the size of a tubulin monomer. A third bound density (bMIP3) had a multi-lobe globular structure bound to the MT lumen wall via three stalk-like densities. A fourth class average (bMIP4) bore similarity to bMIP2, but was bound to the MT lumen with one stalk and harbored an extra density on its top. All bMIPs were found exclusively in the pluripotent P19 cells, with bMIP2 and bMIP1 having the highest and lowest abundances, respectively (**Fig.S6A**).

**Figure 2:**
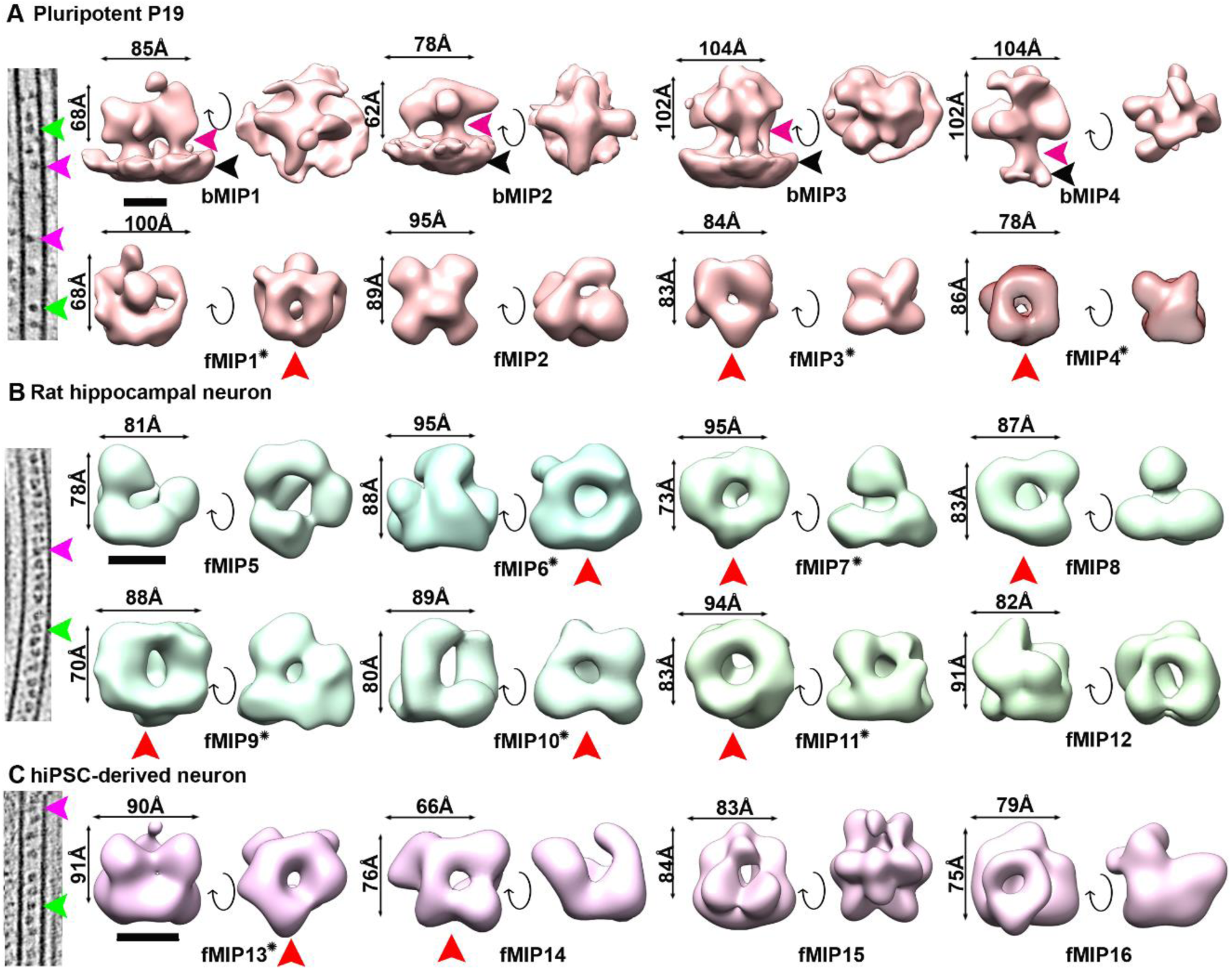
Subtomogram averages of lumenal particles in P19, primary and hiPSC-derived neurons. **A.** Left, a slice of a pluripotent P19 MT showing visually identified bound (magenta arrowhead) and floating (green arrowhead) type of lumenal particles. Class averages (in brown) from the pluripotent P19 cells. MT wall-bound class of particles indicated by their attachment to MT lumen (black arrowhead) with stalk like density (magenta arrowhead). **B**, **C**. Left, same as in (A) for the corresponding neuronal cell-types. Class averages of lumenal particles found in primary neurons (green) and hiPSC-derived neurons (purple). Two-sided arrows indicate maximum dimensions of the maps. Curved arrow indicate 90° rotation. Averages containing ring-shaped scaffold originating from all cell types are indicated by red arrowhead. Cage-like fMIPs with empty cores are marked with a star. Scale bars 5 nm. See also Figures S2, S3 and S4.

The floating particles (fMIP) from all cell types appeared predominantly globular and were located centrally in the MT lumen (**Fig. 2A-C**). STA confirmed that the densities were devoid of any putative stalks. Yet, the absence of a connecting density between the particles and the MT wall does not necessarily preclude a contact, because high flexibility can limit convergence into a discrete density in the average maps. This was suggested to be the case for the absence of a putative stalk density in the MIP average from mouse DRG neurons (red boxed, **Fig. S3**) (21). Visual inspection indicated that most of the fMIPs contain a ring-shaped scaffold – a structural feature common to all the cell-types reported here (**Fig. 1G**) and previously reported for other mammalian neurons (21, 22). STA confirmed that the majority of fMIPs possess a ring-shaped scaffold (**Fig. 2B-C**; red arrowheads), which we further substantiated by analyzing clustering of pair-wise cross-correlation (CC) values between the ring-shaped fMIPs (**Fig.S6B**), implying high degree of similarity at the relatively modest resolution of the averages (22-34Å, **Fig. S4**). A few of the ring-shaped fMIPs exhibited a distinct cage-like topology with an empty core (**Fig. 2A-C;** marked with star: fMIP1, fMIP3-4, fMIP6-7, fMIP9-11, fMIP13). Among them, fMIPs 4 (mouse), 11 (rat) and 13 (human) closely resemble the recently elucidated ring-shaped MIP density in mouse DRG neurons (cc values > 0.8) (21). The finding of ring-shaped scaffold as one of the MIP components in various neuronal cell types underscore their potential general importance in MT biology. However, we found differences among the ring-shaped fMIPs that arise from the configuration and different sizes of the globular densities attached to the ring. Except for fMIP13, angular sampling of all the fMIPs are uniform suggesting these differences are less likely to originate from their orientation bias with respect to the tomographic missing wedge (**Fig. S4A**). Non-ring fMIPs such as fMIP2 (in P19), 5 (primary neurons), and 15 (hiPSCs) are globular and topologically not related to each other. They could therefore be cell-type specific. fMIPs exhibited varied abundances. Among the ring-shaped fMIPs, fMIP1 of pluripotent P19 cells had similar abundance as fMIP 6 and 9 in primary neuron, while fMIP13 of hiPSC-derived neurons had the highest abundance of all (**Fig.S6A**). In addition, unlike all other MIPs analyzed, fMIP13 showed short-range packing with the most frequent NN-distance positioned at 8-10 nm distance within the MT lumen (**Fig.S6C**).

### Taxol treatment alters lumenal particle distribution

In order to identify the molecular constituents of lumenal particles, we sought a way to modulate particle concentrations within the MT lumen to enable a proteomics-based analysis. We hypothesized that Taxol, whose binding is known to occur at the MT lumen (49), could affect MIP distribution. Therefore, pluripotent P19 cells were treated with 5 µM Taxol for 30 mins prior to vitrification and visualized by cryo-ET. Particle concentrations in the Taxol-treated samples were found to be reduced by ∼25% compared to the control (**Fig. 3A; S7A, D; MovieS4**). Further, NN-distance measurement showed that lumenal particles became randomly organized in Taxol-treated cells (**Fig. 3B**) compared to the short-range packing observed in the control cells (**Fig.S2E**). Analysis of the relative abundances of each P19-specific class averages showed while the ratio of bMIPs and fMIPs was close to 1:1, a global reduction in the fMIPs and a significant increase in bMIPs followed in Taxol treated cells (**Fig. 3C, D**). bMIP1 concentration was found to be significantly higher and almost exclusive in Taxol treated P19 cells (**Fig. 3D**), with more particles having short range order with frequent NN-distance of 8 nm compared to the control (**Fig. 3E, 3A**). Therefore, Taxol treatment offered an opportunity to probe the identity of neuronal MIPs.

**Figure 3:**
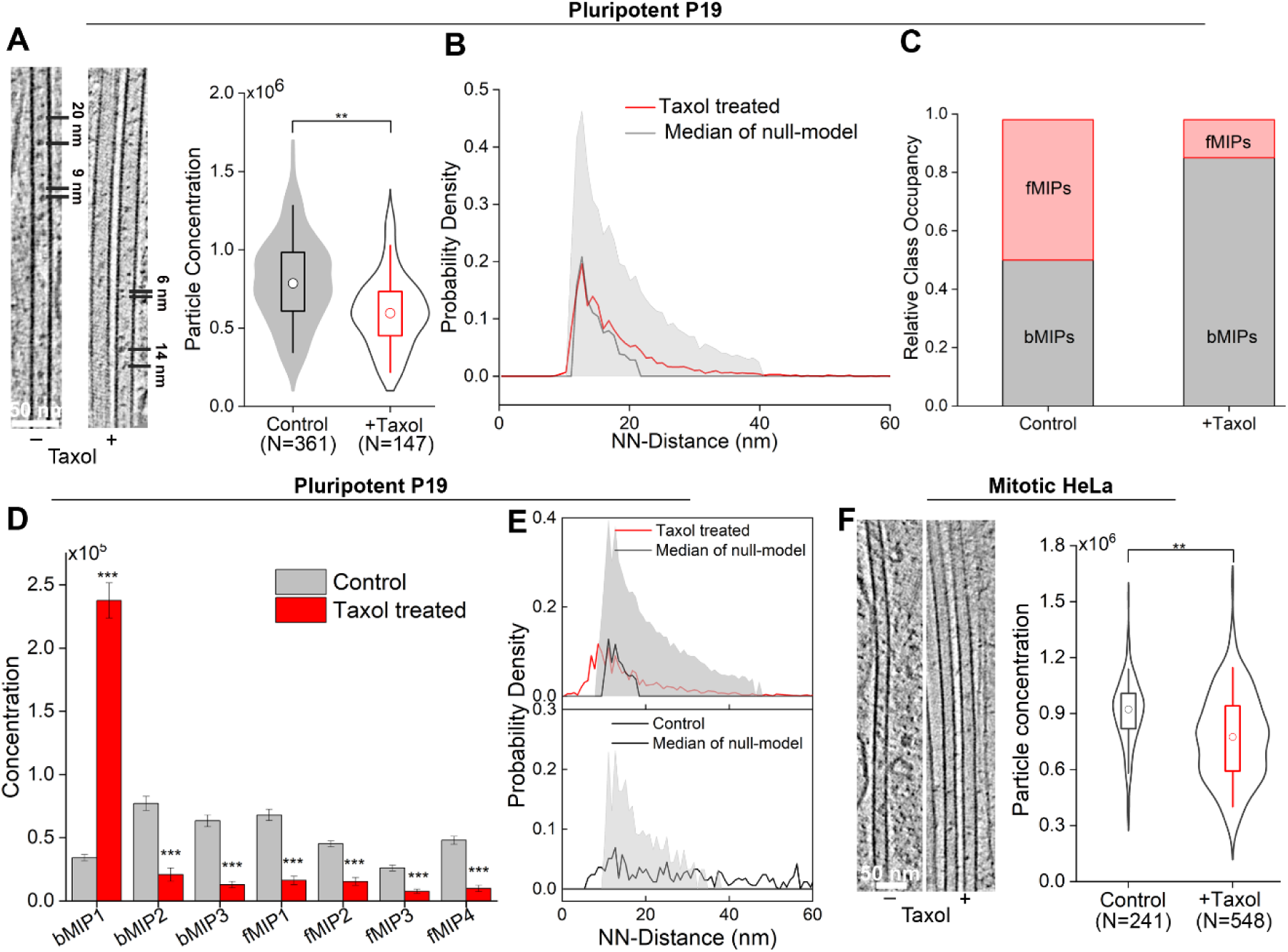
Taxol treatment reduces lumenal particles concentration. **A**. 6.8 nm thick tomographic slices of control (-) and Taxol-treated (+) pluripotent P19 cells showing particle distribution inside MT lumen. Representative distances between the particles are indicated. Quantification of particle concentrations of indicated samples shown as combination of box-plot and data distribution in violin plot. White circle denotes median. Statistical significance using Mann-Whitney test, **p<0.01. N, number of MTs. **B**. NN-distance distributions of the lumenal particles in indicated condition. Shaded gray area indicate IC [5, 95] %. **C**. Relative abundance of wall-bound (bMIPs) and floating (fMIPs) category of class averages in control and Taxol-treated pluripotent P19 samples. **D**. Concentrations of the class averages in control and Taxol treated pluripotent P19 cells. Error bar indicates standard error of mean (SEM). Statistical significance ***p<0.001, obtained by two-sample t-test. **E**. NN-distance distribution of bMIP1 in control and Taxol-treated pluripotent P19 cells. Shaded gray area indicate IC [5, 95] %. **F**. Representative 9 nm thick tomographic slices of control and Taxol-treated mitotic HeLa cells showing particle distribution inside MT lumen. Quantification of particle concentrations shown as in (A). Circle in the middle denotes median. Statistical significance obtained using Mann-Whitney test, **p<0.01. See also Figure S7.

### A subset of fMIPs identified as putative tubulin binding cofactors

While the mechanism by which Taxol leads to reduction of lumenal particles abundance, especially of fMIPs, is not clear, we leveraged this observation as an experimental condition to identify Taxol-sensitive MT-associated proteins using a differential enrichment of tubulin interacting proteins in control and Taxol-treated cells by affinity purification-MS (AP-MS) based proteomics. Towards that end, we opted to use non-neuronal HeLa Kyoto cells that offered a technical advantage as they can be grown in sufficient quantities suitable for pulldowns. Mitotic HeLa (in contrast to interphase cells) expressing GFP tagged β-tubulin (26, 50) exhibited lumenal particles with similar abundances as in pluripotent P19 cells (**SI Appendix, Fig. 3F; Fig. S7B-C, S7E-G**). Taxol treatment similarly reduced the abundance of lumenal particles in the mitotic HeLa (**Fig. 3F**). Visual inspection showed that the HeLa lumenal particles share morphological similarities, including the ring-shaped particles, with those of the neuronal cells (**Fig. S7H**). Therefore, β-tubulin was isolated by affinity purification using anti-GFP antibody immobilized on magnetic beads (**Fig. S8A**) from intracellularly crosslinked control (DMSO-treated) and Taxol-treated mitotic HeLa cells (**SI Appendix**). Cross-linking ensured preservation of information on transient MT-interacting particles during the isolation procedure. Differentially enriched proteins were identified by mass spectrometry (**Fig. S8B, C**). We found that Taxol treatment significantly reduced the abundance of over 160 cytoskeleton-related proteins, including several structural MAPs known to bind the MT outer surface, such as Ensconsin, MAP4 and MAP1S (see Data availability, **TableS2, Fig. S8B, C**). Notably, we observed a modest reduction of three cytosolic tubulin binding cofactors (TBC): Tubulin binding cofactor A (TBCA), D (TBCD) and E (TBCE) (P <0.05) (**Fig. S8C**).

We next systematically analyzed all the hits belonging to the Gene Ontology (GO) term of MT cytoskeleton-related proteins by examining structures (if available) or structural predictions (51) of them. All models and available structures were low-pass filtered to the resolution of the density maps generated for the lumenal particles for comparison (**Table S2**). Our analysis showed that most *bona fide* MAPs found in this study including the neuron-specific MAP6 that is suggested to be a component of lumenal particles (42), are predicted to be disordered and therefore unlikely to correspond to our globular density maps (**Fig. S9, Table S2**) (21). Notably, we did not find TAT, the most commonly perceived component of lumenal particles in our AP-MS hits even though mitotic MTs are known to be highly acetylated, similar to neuronal MTs (52). Interestingly, ring-shaped scaffolds of the cage-like fMIPs (**Fig.2A-C**, marked with star) were found to bear high similarity (with a cross correlation (cc) of >0.8) to the 24 Å negative stain EM-derived map of tubulin binding cofactor-D (TBCD, mass 132 kDa, EMD-6390/6392, **Fig. S9**) (53), involved in folding and degradation of β-tubulin (**Fig. 4A**)(54). We used the crystal structure of the yeast Cse1p (PDB:1Z3H) (55) that structurally mimics TBCD (53), filtered to 25 Å resolution to fit into our ring-shaped EM density using rigid body fitting protocol (**SI Appendix**). The distinct alpha-solenoid ring structure of the Cse1p made up with HEAT repeats fitted the ring-shaped base of the fMIP1, 9 and 13 (cc > 0.8, green, **Fig**. **4B-D**). Previous biochemical evidences indicate that TBCD can exist in a stable complex with ADP ribosylation factor-like protein 2 (Arl2) (56) and with β-tubulin (57). In case of fMIP1, Arl2 model (PDB: 1KSH) fitted well (cc 0.95) within the globular density (light pink), suggesting a complex between TBCD and Arl2 (**Fig**. **4B**) with a minimum projected mass of ∼150 kDa. A portion of the density (orange) remained unassigned. A β-tubulin density fitted in the remaining globular density of fMIP9 with a correlation coefficient of 0.85 (yellow, **Fig**. **4C**). Therefore, fMIP9 tentatively represents a binary complex between TBCD and β-tubulin with a projected mass of ∼180 kDa. In case of fMIP13, portions of the density (grey and magenta) remain unassigned (**Fig**. **4D**). The mass of these complexes correlates with our mass measurements from the tomograms (**SI Appendix, Fig.S5F**). The U-shaped density with two globular heads of fMIP5 (cyan, **Fig**. **4E**) bore structural resemblance to the negative stain EM-derived map of TBCE (EMD-2447), another cofactor that binds α-tubulin (58). Therefore, as in the TBCE density map (EMD-2447) the globular heads of fMIP5 fitted with the Cap-Gly (PDB: 1WHG) and UBL domains (PDB: 4ICV) while the curved segment between the globular heads likely corresponds to the TBCE LRR domain (58). The curved segment fitted TLR4 (53) that structurally mimics the TBCE LRR domain (cc 0.94, PDB: 3RJ0). The Cap-Gly domain binds to a globular density in our map that can accommodate α-tubulin (yellow, with cc 0.88, **Fig**. **4E**, PDB: 1TUB), and importantly places the tubulin binding loop close to the tubulin C-terminus. Based on the above assignments, we suggest putative identities for 9 ring-shaped lumenal particles (fMIP1, fMIP3-4, fMIP6-7, fMIP9-11 and fMIP13) and one non-ring shaped (fMIP5) out of the 16 fMIPs to be TBCs bound to tubulin. These accounted for a high fraction of MT lumenal particles in the different neuronal cell-types (**Fig. S6A**, **Table S1,** up to 75% of all fMIPs in primary neuron).

**Figure 4:**
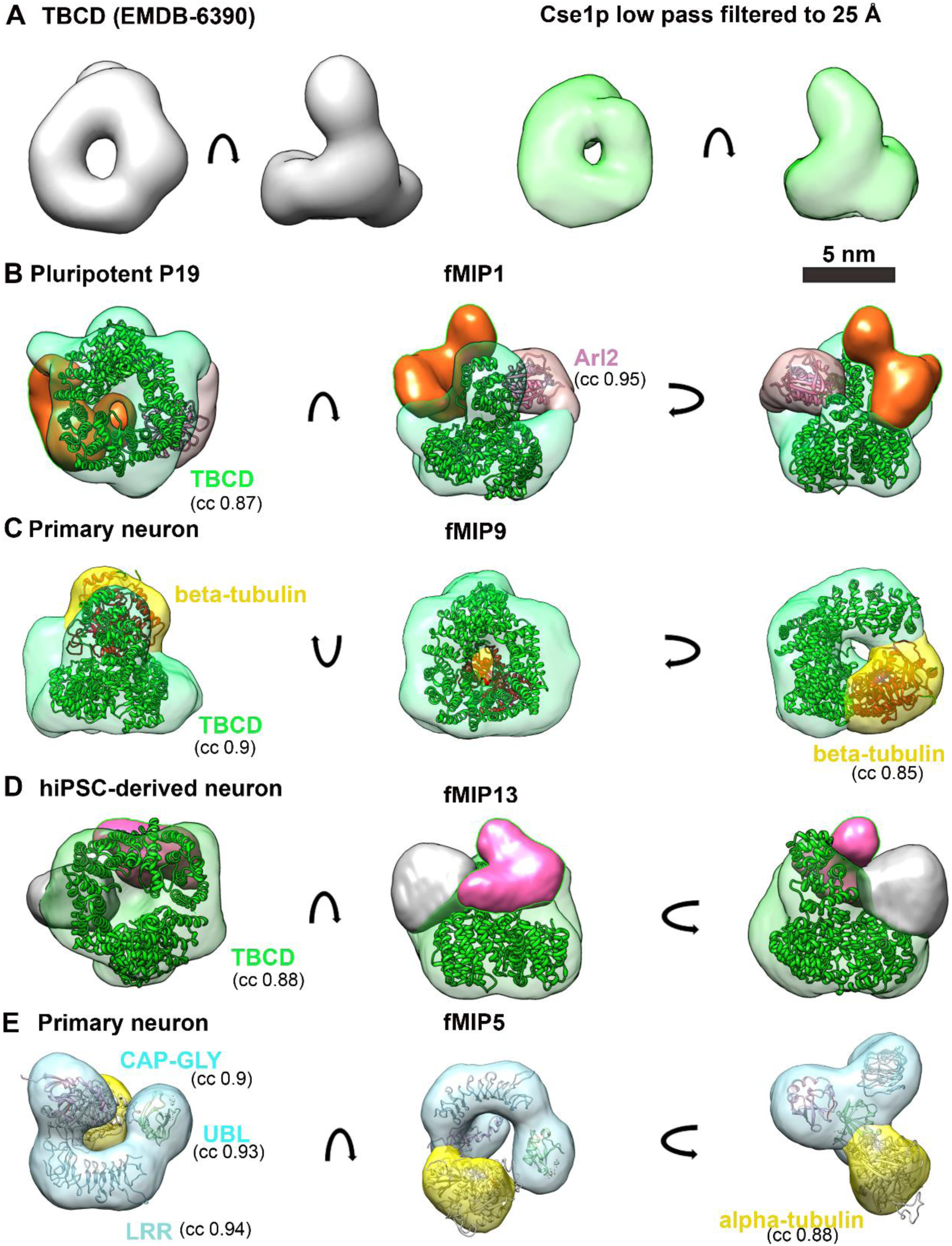
A subset of fMIPs resemble tubulin-binding cofactors. **A**. Single particle EM map of TBCD obtained from EMDB-6390 (53) showing ring-shaped base. For comparison, low pass filtered map of Cse1p (PDB: 1Z3H) (55) at 25 Å is shown in two views. **B**. Segmented density map of fMIP1 from pluripotent P19 cells shown in three views. The ring-shaped base (green) is fitted with HEAT repeats of Cse1p (green). Light pink globular density is fitted with the GTPase, Arl2 (PDB: 1KSH). Remaining density (orange) is unassigned. **C**. Segmented map of fMIP9 from primary neurons shown in three views. The ring-shaped base (green) is fitted with HEAT repeats of Cse1p TBCD (green). Yellow globular density is fitted with β-tubulin (PDB: 1TUB) as per the biochemical evidence. **D**. Segmented map of fMIP13 from hiPSC-derived neurons shown in three views. The ring-shaped base (green) is fitted with HEAT repeats of Cse1p (green). Remaining densities (gray and pink) are unassigned. **E**. Segmented map of fMIP5 from primary neurons shown in rotated views. The U-shaped base (cyan) is fitted with LRR (PDB: 3RJ0), and two globular head domains are fitted with UBL (PDB: 4ICV) and CAP-Gly (PDB: 1WHG) domains of TBCE as indicated. Yellow globular density is fitted with α-tubulin. CC values from rigid body fitting of each model low pass filtered to 25 Å are indicated. See also Figure S6 and S9.

### MT lumenal particle abundance increases in neuronal differentiation

Observations of densely and periodically decorated MT lumens particularly in rodent and hiPSC-derived neurons (**Fig.1A-B, D**), as well as in other mammalian and insect neurons (17, 21–24), raise a question as to whether such decoration represents a characteristic of differentiated neuronal MTs. To address this question, we took advantage of the ability of pluripotent P19 cells to differentiate and develop neuronal processes *in vitro* (59). Neurospheres generated after aggregation of the P19 cells upon exposure to 0.5 µM retinoic acid (RA) were allowed to differentiate for about 1 week. Neuron-like cells were selectively grown using a DNA synthesis inhibitor (**SI Appendix**). When these *in vitro* differentiated neurons were imaged using cryo-ET, their thin processes revealed parallel arrays of MTs (**Fig**. **5A-C**). Unlike pluripotent P19 cells, the MT were frequently observed embedded in a dense intermediate filaments network (**Fig. 5B**) (59, 60). The cryo-ET data revealed significantly higher concentration of tightly packed particles relative to their parental state, arranged in a periodic array with 8-10 nm NN-distance (**Fig. 5D-E**). Thus, emergence of dense and ordered arrays of lumenal particles occurs after *in vitro* neuronal differentiation and could be a characteristic of neuronal MTs (17, 21–24). STA further showed enrichment of only P19-speciifc particles in the MT lumen after differentiation, with a significant increase in the concentrations of bMIP4 and fMIP4 (**Fig. 5F**). In particular, lumenal particles with an ability to bind to the MT wall (such as bMIP4) and/or invoke tubulin proteostasis (fMIP4) could reflect emergent properties of the MT cytoskeleton in adaptation to neuron-specific functions.

**Figure 5:**
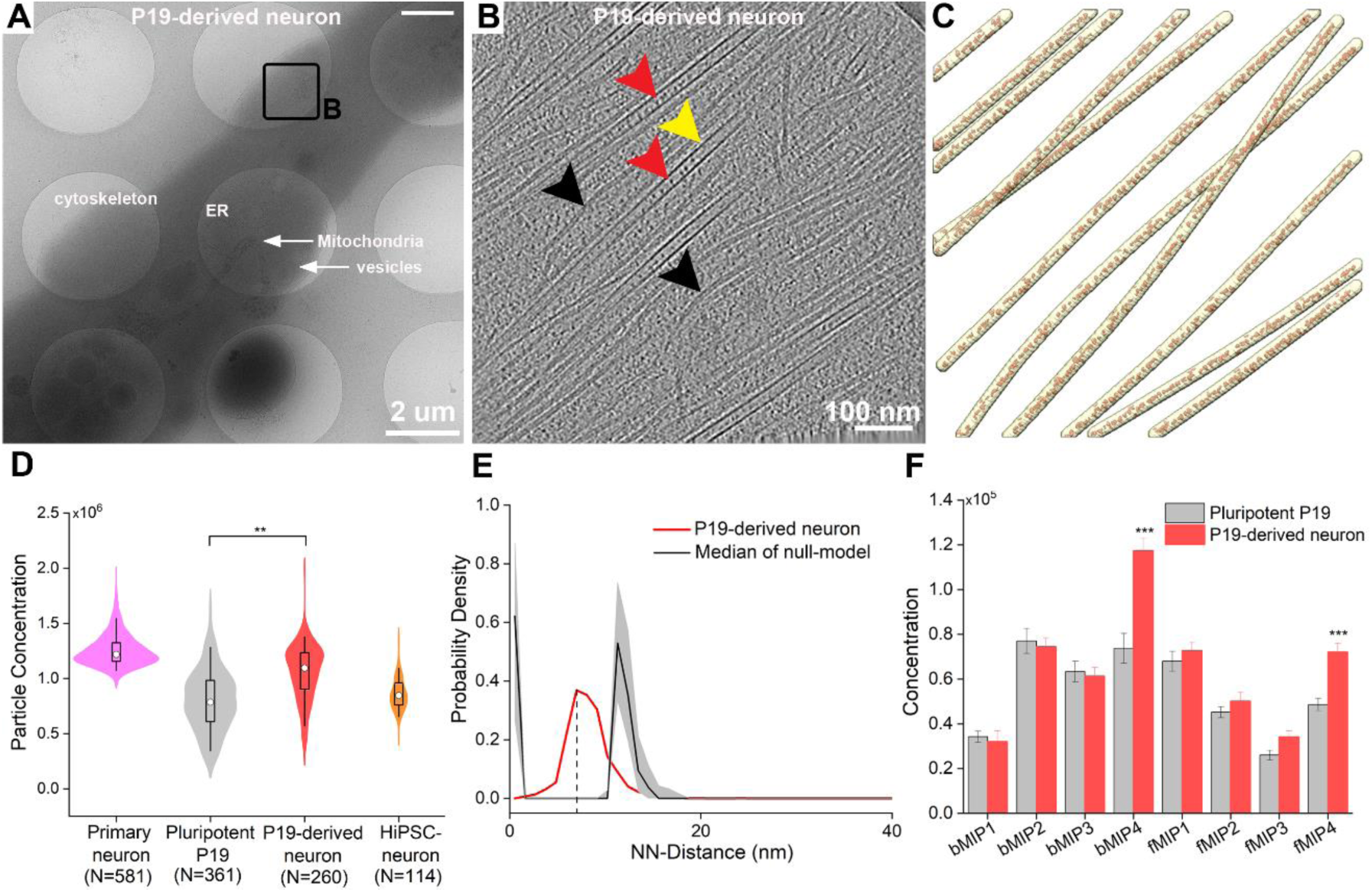
Induction of neuronal differentiation enhances MT lumenal particles. **A**. A P19-derived neuronal process at 10 days *in vitro* differentiation on EM grid, showing dense cellular organelles and a polarized cytoskeleton. **B**. 9 nm thick tomographic slice of a P19-derived neuronal process showing MTs (yellow arrowhead) containing stacked lumenal particles (red arrowheads), and parallel array of Intermediate filaments (black arrowheads). **C**. 3D rendering of traced MT filaments. MTs (yellow), lumenal particles (red). **D.** Quantification of the particle concentrations (numbers/µm^3^ of lumenal volume) represented as boxplot within a violin. For comparison, concentrations for primary and hiPSC-derived neuron from Fig.1I are replotted here. Median values are marked by a circle. Asterisks indicate Mann-Whitney test significance: **p<0.01. N, number of MTs. **E**. NN-distance distributions of the lumenal particles in P19-derived neurons. Black dashed line indicate most represented NN-distance. **F**. Comparison between the concentrations of each class averages before and after differentiation. Error bar indicates SEM. Statistical significance ***p<0.001, obtained by two-sample t-test.

### Lumenal particle abundance correlates with MT curvature, lattice defects and freshly polymerized plus ends

We wondered whether the enrichment of lumenal particles in differentiated cells is linked to measurable MT properties. Cryo-tomograms revealed curved MTs in neuronal processes, with segments exhibiting sinusoidal trajectories (**Fig. 6A**). We first quantified the extent of MT curvature using the tangent-correlation length (^a^L_p_, **SI Appendix**) (26). ^a^L_p_ in both primary and hiPSC-derived neurons indicated high curvature with mean values of 28.4±3.4 µm and 21.7±4.2 µm (**Fig.S10**). In contrast, pluripotent P19 MTs were less curved with a broad ^a^L_p_ distribution and a mean of 41.6±5.33 µm (**Fig. 6C, Fig.S10**). Upon P19 differentiation *in vitro*, the distribution narrowed and centered on 25.9±3.9 µm, at par with the curvatures found in primary neurons (**Fig.S10**). Examining the correlation between MT curvatures and lumenal particle concentrations showed that in primary neurons, highly curved MTs contained higher particle concentrations (**Fig. 6B**). A similar trend was observed in hiPSC-derived neurons, for which only 4 data points could be measured. In pluripotent P19 cells, a distinct segregation into two populations was observed where highly curved MTs contained high particle concentrations (similar to primary neurons) (**Fig. 6C**) and vice versa. Upon differentiation, the P19 cells developed a trend similar to the primary neurons (**Fig. 6D**). This correlation suggested that lumenal particles, predominantly bMIPs including bMIP4, preferentially localized within curved MTs, potentially contributing to stabilization of curved MT lattice in a manner similar to that suggested for lumenal MAP6 (42).

**Figure 6:**
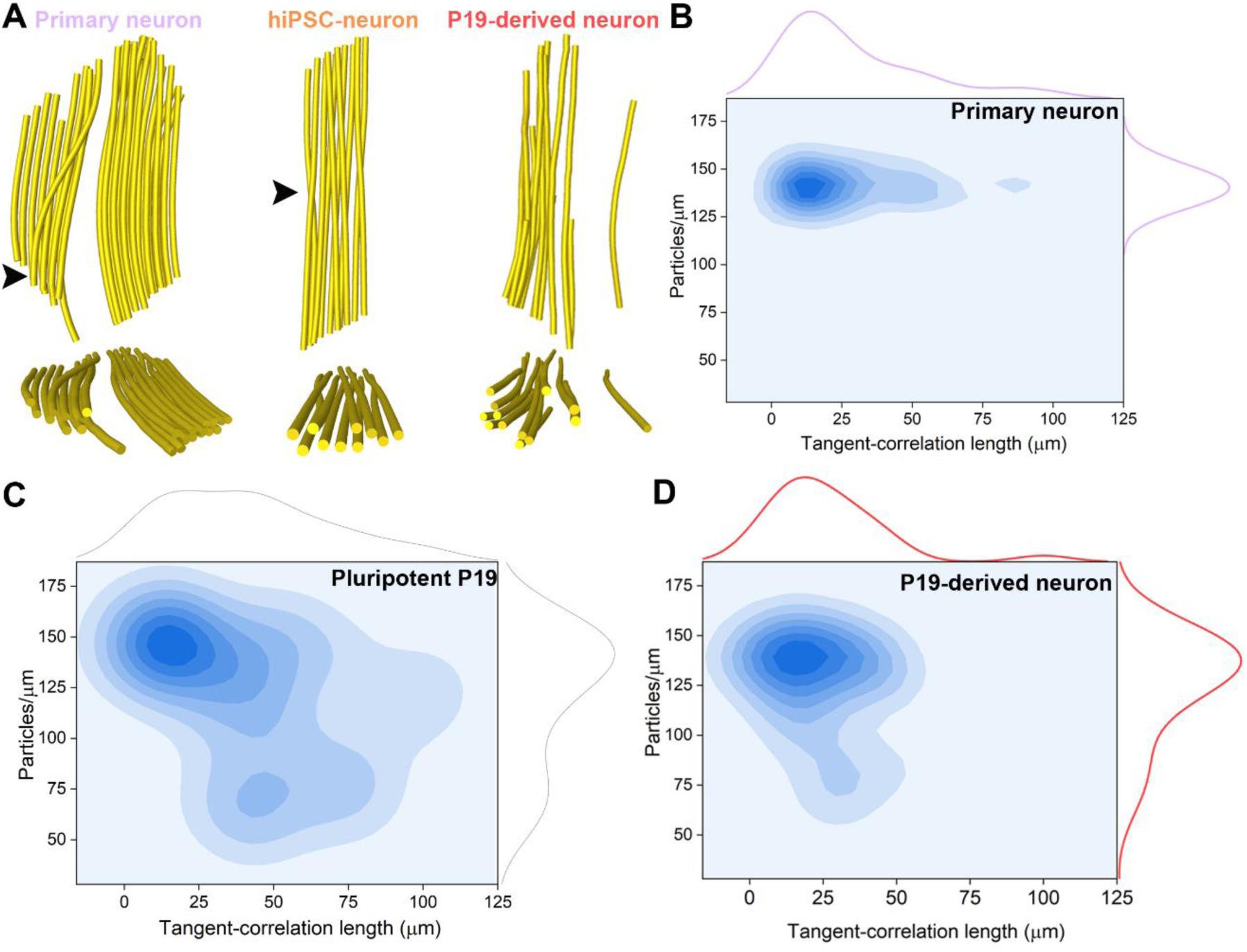
MT curvature correlates with lumenal particle concentration. **A.** Representative 3D rendering of MT bundles from a primary, hiPSC and P19-derived neuronal process. Top, x-y views, bottom, x-z views. Black arrowheads indicate MTs displaying curved trajectory. **B-D.** Distribution plots of tangent-correlation lengths (^a^L_p_) and mean particles per micron for **B**) primary neurons (magenta) (N=40), **C)** pluripotent P19 cells (gray), (N=24), and **D**) P19-derived neurons (red) (N=26), shown on the margins. Correlation between the ^a^L_p_ and mean particles per micron indicated as contours for the indicated cell types. N denotes numbers of tomograms analyzed. See also Figure S10.

High MT curvature could lead to breaks (26), which may represent entry points into the lumen, as was suggested for TATs (61). We therefore next examined the abundance of lumenal particles with respect to partially broken and exposed MT lattice regions in pluripotent P19 cells. We pinpointed several partially broken MTs showing loss of one or several protofilament segments (**Fig. 7A**). A ‘lattice map’ of one such representative case using STA, where individual subtomograms were mapped back to the original positions in the tomogram and color coded according to the cross-correlation coefficient (CCC) (**Fig. 7B-C**), showed that CCC-values flanking the broken site were lower, in accordance with a disordered lattice. We found that the pF number in the flanking regions of the severed lattice was 13, in contrast to the observed pF heterogeneity near lattice defects *in vitro* (**Fig. 7C**) (62). Lumenal particle in these regions were counted and represented as a color coded spatial occupancy map. Such representation highlighted the presence of a ‘micro-cluster’ flanking the broken region compared to the intact part (**Fig. 7D**, red segment). We used bivariate Ripley’s L function to derive statistical significance of such clustering (46). Distances between the particles and break points were measured in 15 such instances and compared with complete randomness using Monte Carlo simulation. The experimental median of the L function displayed a bell shaped curve with positive values (green line, **Fig. 7E**) that is distinct from the random simulation (gray area, **Fig. 7E**), indicating clusters of a size of ∼300 nm around the lattice breaks were statistically significant (**Fig. 7E**).

**Figure 7:**
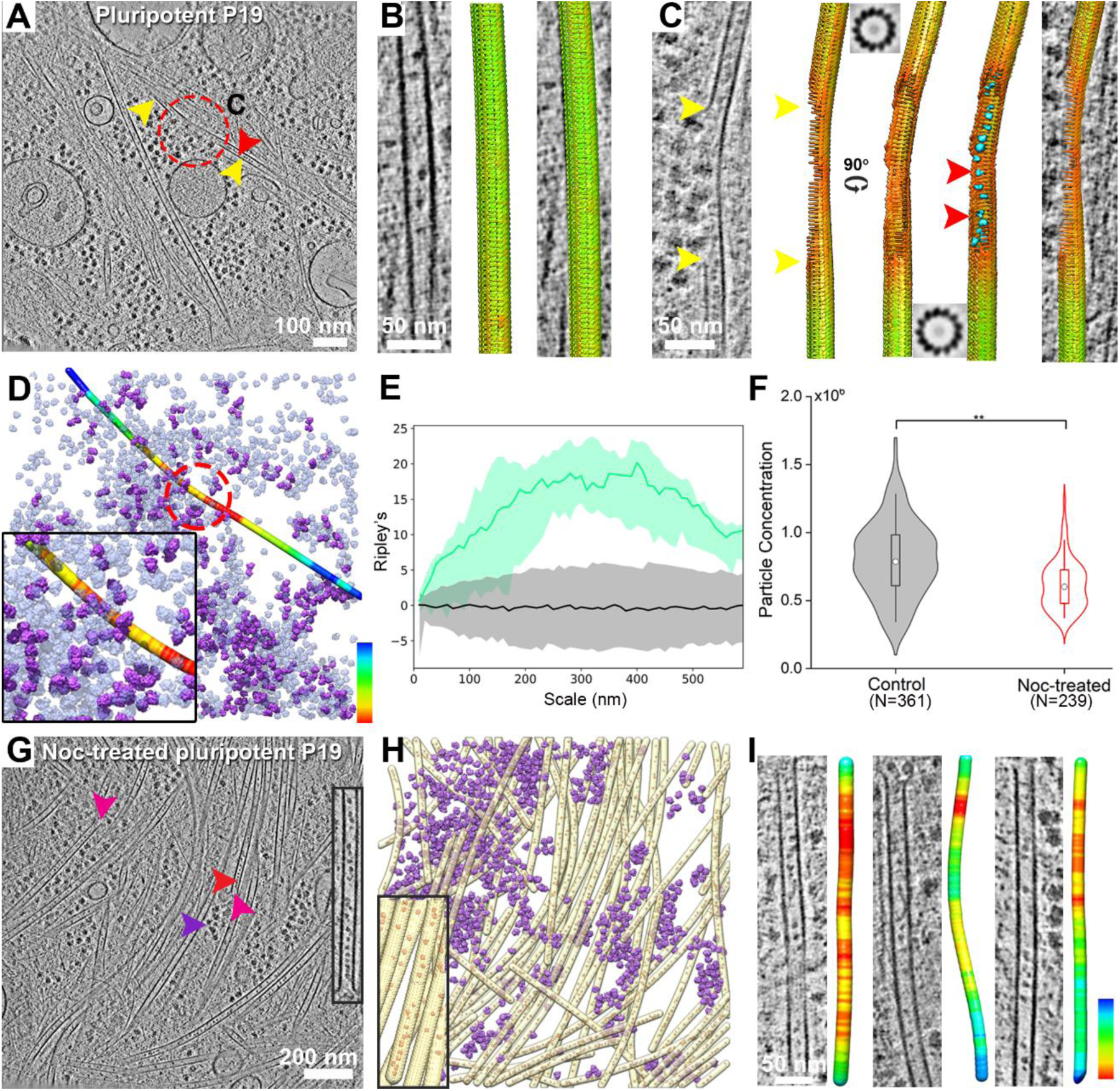
Lumenal particles cluster at the lattice breaks and MT plus ends. **A.** 6.8 nm thick tomographic slice of a pluripotent P19 MTs showing a broken MT (yellow arrowheads and red circle, enlarged in C). Red arrowhead indicates lumenal densities. **B**. 6.8 nm thick tomographic slice of an intact pluripotent P19 MT along with lattice map superimposed. Subtomograms are represented as arrows and color coded by CCC values where green represent highest CCC. **C**. Similar to B, but for a partially broken MT lattice (between yellow arrowheads). Lattice maps shows missing subtomograms in the damaged regions. 3D-surface rendered lumenal particles (cyan) are mapped back in the exposed MT region. STA confirmed 13 pFs in the flanking regions of severed lattice (2D cross sections of averages at top and bottom). **D**. Surface rendered image of the tomogram in A, with the single MT color coded with lumenal particle occupancy where red represent highest abundance. Ribosomes shown as transparent violet. Ribosomes that form polyribosome shown in opaque violet (Fig. S5G-I). Inset is a zoomed image of the interface (red circle). **E**. Bivariate Ripley’s L function for the particles clusters found at the partially broken MT lattices (N=15 MT breakpoints) (green line). Black line indicate median of the L function for a set of random particle distributions and the grey shaded area represent the IC [5, 95]%. **F**. Quantification of particle concentrations in control and freshly polymerized MTs of pluripotent P19 cells. Statistical significance by Mann-Whitney test **p<0.01; N, number of MTs. **G**. 6.8 nm thick tomographic slice showing particle distribution inside freshly polymerized MT lumen. Arrowheads indicate open-end of MTs (magenta), lumenal particles (red), and ribosomes (violet). Inset shows a zoomed image of a MT open-end. **H**. 3D surface rendering of panel G showing the dense MT network (yellow) along with lumenal particles (red) interspersed with ribosomes (violet). Inset is an enlarged region of MT open-end from (H). **I**. Representative slices of plus-end (top side) of freshly polymerized MTs of pluripotent P19 cells with corresponding color coded occupancy map of lumenal particles along the MT length (red represent highest abundance). See also Figure S7 and S11.

We next tested whether micro-clustering also occurs at the open MT ends representing another possible entry point for lumenal particles. To this end, MT polymerization was initiated in pluripotent P19 cells by a brief Nocodazole treatment and washout (**SI Appendix**) (26). While particle concentration was overall lower in newly polymerized MTs compared to the control (**Fig. 7F**), lumenal particle clustered near the MT open-ends (**Fig. 7G-I, MovieS5**). We statistically evaluated such clustering using the previously described bivariate Ripley’s L function, which revealed a weak tendency of the particles to cluster with domain size up to 200 nm from the MT open-end (see **Fig. S11A-B** for details). The particles were more tightly packed with a mean inter-particle distance of 9.6±3.4 nm within ∼100 nm from the MT open-end compared to distal regions (mean distance 14.3±6.7 nm) (**Fig. S11C**). STA showed that freshly polymerized MTs accumulated similar classes of both bMIPs and fMIPs as the control P19; among the class averages, concentrations of bMIP3, fMIP1, 2 and 4 were significantly reduced (**Fig.S11D**). Visual inspection showed that both bMIPs and fMIPs are part of the micro-cluster observed near the MT open end. Our data thus suggests that particles can access the lumenal space in small clusters at MT lattice breaks or via the open-end of the polymerizing MTs. The presence of such clusters might impact nascent MT-tip during polymerization or sites of lattice damage through tubulin lattice stabilization.

## Discussion

The presence of lumenal particles inside MTs has been known since decades (13–17, 63). Yet, our understanding of the molecular components contributing to the formation of these particles and their functions remained elusive. It is an open question whether MT lumenal particles represent novel types of MAPs involved in modulating MT stability, or whether they could be proteins or mRNA that are stored or transported inside MTs (23, 24). We leveraged *in situ* cryo-ET combined with STA and targeted proteomics to extract structural, spatial and biological information about MT lumenal particles of various neuronal cells.

The periodic decoration of the MT lumen by particles appears to be a key feature of neuronal MTs **(Fig. S2D-F, Fig. 5E)** (17, 21–24). Such decorations could endow properties such as stability to neuronal MTs (64). Extensive mechanical forces are expected to be exerted on MTs during neuronal development (65), trafficking (2), and axon migration (66). As a force-bearing element, neuronal MTs are subject to bending and coiling, resulting in highly curved appearance *in situ* (26, 42) (**Fig. S10**). The densities we resolve for bMIPs (**Fig. 2A**) are well placed to structurally stabilize curved MT lattices by crosslinking several pFs together via multiple stalk-like densities (positioned 4 nm apart), in analogy to MIPs related to MT doublets in motile cilia and flagella that function to stabilize the MT lattice (32). For cytoplasmic MTs, MAP6, an intrinsically disordered protein with three MT binding Mn modules is known to stabilize coiled neuronal MT lattice in a similar manner from the lumenal side (42). However, its predicted disordered structure precludes it from representing any of the bMIP densities (**Fig. S9**) (21). The emergence of curved MTs during the course of neuronal differentiation of P19 cells with concomitant enrichment of the bMIPs further supports this hypothesis. Furthermore, in the pluripotent P19 cells, higher concentration of lumenal particles was correlated with more curved MTs and vice versa. Therefore, bMIP localization or enrichment seems to happen selectively to those MTs that experience larger mechanical forces, and possibly suggest their protective role in maintaining MT lattice architecture. Taken together, bMIPs may add another layer to the existing mechanisms that regulate material properties of neuronal MT along with structural MAPs and a complex tubulin-code (67).

Our proteomics data combined with structural modeling suggests that the cage-like fMIPs containing a ring-shaped scaffold with an empty core could be TBCs bound to monomeric tubulin (**Fig. 4B-E**). TBCs are ubiquitously found in all cells and are essential for ‘tubulin proteostasis’, including folding of the tubulin monomers, formation of tubulin heterodimers and for tubulin degradation (68). Thus, the presence of TBC-bound tubulin complexes within the lumen could have interesting implications. Currently, not much is known about tubulin biogenesis or the localization of its folding machinery in neuronal cells, especially in axons (69). Tubulin must be present in sufficient amounts in the distal regions of axons to support nucleation and dynamics of MTs for the maintenance of synapses and growth.

However, axonal processes contain low numbers of ribosomes, in agreement with low protein synthesis rates along axons (70). Furthermore, tubulins are transported from the cell body to distant neurites/axons at a slow speed of 0.1-3 mm/day (71). Therefore, it is interesting to consider a mechanism wherein TBC-bound tubulins packed within the MT lumen could locally provide fresh dimers for the incorporation to lattice defects in distal regions of neurons. In growth cones or synapses where local translation is active, fMIPs carried by the MTs could also coordinate with translating ribosomes to replenish local tubulin pool essential for new MT nucleation and growth required for growth-cone expansion or synaptic stability. On the other hand, MT disruption by bare TBCD is known to be regulated by Arl2 (72). Therefore the presence of TBCD:Arl2 complex might ensure maintenance of MT cytoskeletal integrity by removing damaged tubulin from the lattice (56). Both TBCE and TBCD could also take part in recycling of misfolded tubulins that pop out of the lattice due to MT bending, along with TBCB and TBCA, respectively. It should be noted that we have not detected TBCB (27 kDa) and/or TBCA (13 kDa) in our class averages possibly due to their small sizes and the fact that they are mainly cytosolic working immediately downstream of CCT chaperonin that folds tubulins. Taken together, fMIPs could maintain a local tubulin pool and provide fresh tubulin dimers for MT repair and growing MTs required for growth cone expansion, axon branching and maintenance of synapses, or for repair of damage along the MT lattice (73).

Long-lived neuronal MTs are suggested to accumulate nanoscale lattice defects due to extensive synaptic trafficking and force-bearing during neuronal growth (25, 26, 74). Generally, depolymerization of defective MTs, either due to their intrinsic property or by the action of severing enzymes, and polymerization of a new set of MTs has been regarded as the main mechanism for maintaining a functional MT network. However, such wholesale replacement mechanism in long neuronal processes could disrupt neuronal function. In such scenario, bMIPs could stabilize the damaged MT lattice by forming micro-cluster around the defect and holding the surviving pFs together. Lattice defects might act as a good substrate for MT-severing enzymes such as Katanin (75). bMIPs clusters could potentially antagonize severing enzyme function and provide sufficient time for MT self-repair through fresh tubulin dimer incorporation. A similar protective role has been recently proposed for the CLASPs that localize to bent MT regions and stabilize them (76). At the same time, fMIPs could invoke local tubulin homeostasis at sites of lattice defects, provide fresh tubulins for their incorporation in the damaged lattice and remove damaged tubulins. This process presents an optimal way to self-repair while maintaining the complex ‘tubulin code’ required for long lived neuronal MTs (73). Our findings could further shed light on the potential role of impaired tubulin proteostasis resulting in defective MT networks observed in severe neuro-developmental disorders such as giant axonal neuropathy (77), hyperparathyroidism (78), inherited early-onset encephalopathy (79) and infantile neurodegeneration (80).

In summary, we elucidate the native organization and morphology of neuronal MT lumenal protein complexes and suggest that a subset represents a number of tubulin-binding chaperone complexes. We hypothesize that the MT lumen could serve as a transport channel for delivering fresh tubulins (23, 24) that play a role in maintaining MT lattice integrity under compressive or transport-related forces in neurons by invoking local tubulin homeostasis, or that provide the essential building blocks for MT growth at distal end of neuronal processes.

## Materials and Methods

### Cell culture and grid preparation

Rat hippocampal neurons derived from embryonic rats (E17-21), murine pluripotent P19 cells, hiPSC-derived neurons and HeLa cells were cultured in appropriate cell culture media in dishes containing pre-treated EM grids according to methods described in the **SI Appendix**. Grids were vitrified using a Vitrobot Mark 4 at 37°C. Please see **SI Appendix** for details.

### Cryo-TEM and tomography

Tilt series for all the cells were collected on a Titan Krios (Thermo Fisher Scientific) operated at 300 kV equipped with a field-emission gun, a Quantum post-column energy filter (Gatan) operated in the zero-loss mode using a 20 eV slit, a K2/K3 Summit direct detector camera (Gatan) operated in counting mode, and a Volta phase plate (VPP) (Thermo Fisher Scientific) for part of the data. Data were recorded with SerialEM software (81) with dose-symmetric (82) or asymmetric bi-directional tilt scheme as detailed **SI Appendix**. Only the best quality tomograms with good contrast produced from cellular areas thinner than 200 nm were used for further analysis (**Table S1**).

### MT tracing and curvature measurement

MT segmentations were performed with the automated filament tracing module in Amira software v.6.2.0 (Thermo Fisher Scientific) as described before (26). The coordinates of the traced filaments were resampled in MATLAB (MathWorks) to obtain equidistant points. MT curvature was determined by measuring the tangent-correlation length as described in our previous work (26). Please see detailed method in **SI Appendix**.

### Density tracing and particle picking

For density tracing and particle picking, tomograms were smoothed by Gaussian low-pass filtering at σv=0.5 pixels and MT lumen segmented based on the MT centerline tracing. PySeg (46) was used to pick particles in a template-free manner as detailed in the **SI Appendix**.

### Subtomogram averaging

Subtomogram analysis for lumenal particles was performed using RELION (version 3.0.5) following published protocols (83). During the refinement, particle half-datasets were processed independently. Unless stated explicitly, all refinements were performed *de novo*, that is, without the use of external references. Datasets corresponding to different cells were processed separately. Extensive classifications were performed to derive maps of different particle populations. Please see **SI Appendix** for details.

### Statistical analysis

Statistical analyses were performed with Origin 9.2 (OriginLab). All boxes in the box-plot representations are bound by the 25th–75th percentile, whiskers span 5th to 95th percentile. Mean values were marked by a black box. Box plots are embedded inside a violin which indicate distribution of the data. For statistical significance, nonparametric Mann-Whitney tests were performed unless stated otherwise. Significances were indicated by stars where ** indicated P<0.01.

## Acknowledgments

We thank I. Poser and A. Hyman for providing the HeLa cell lines, C. Papantonio and C. Capitanio from the Baumeister group for providing rodent primary neuron culture, the MPI Biochemistry mass spectrometry core facility for proteomics support, and A. Petrovic for valuable suggestions regarding homology modeling. S.C. was supported by a Max Planck Society fellowship. A.M.-S. was supported by the Deutsche Forschungsgemeinschaft (DFG, German Research Foundation) under Germany’s Excellence Strategy - EXC 2067/1-390729940. M.T-N and I.H were supported by a fellowship from the EMBL Interdisciplinary Postdoctoral Program (EI3POD) under Marie Skłodowska-Curie Actions COFUND. W.B. acknowledges support from the Center for Integrated Protein Science, Munich. J.M. acknowledges funding from the EMBL.

## Author Contributions

S.C., W.B. and J.M. designed the research. S.C. performed sample preparations, cell biological and biochemical analyses, mass spectrometry based proteomics, cryo-ET on P19 and rodent primary neuron and subtomogram averaging for all data; J.M. provided HeLa tomograms; A.M.-S. developed automated particle picking procedure, spatial statistical tools and helped with data analysis; F.B. provided computational support and helped with data analysis; M.T-N. and J.M. collected hiPSC cryo-ET data; KM.N. and I.Y.H. cultured hiPSC and performed differentiation with M.T-N.; S.C. J.M. and W.B. wrote the manuscript with contribution from the all authors.

## DATA AVAILABILITY

Tomograms included in this manuscript and all maps generated in this work have been deposited in the EMDB, with the following accession numbers: Primary neuron tomogram, EMD-15474; hiPSC-derived neuron tomogram, EMD-15437; Pluripotent tomogram P19, EMD-15440; Differentiated P19 tomogram, EMD-15438; Noc-treated P19 tomogram, EMD-15442; Taxol-treated P19 tomogram, EMD-15443; bMIP1, EMD-15453; bMIP2, EMD-15454; bMIP3, EMD-15455; bMIP4, EMD-15456; fMIP1, EMD-15441; fMIP2, EMD-15439; fMIP3, EMD-15436; fMIP4, EMD-15444; fMIP5, EMD-15457; fMIP6, EMD-15458; fMIP7, EMD-15459; fMIP8, EMD-15472; fMIP9, EMD-15460; fMIP10, EMD-15461; fMIP11, EMD-15462; fMIP12, EMD-15463; fMIP13, EMD-15464; fMIP14, EMD-15466; fMIP15, EMD-15467; fMIP16, EMD-15468; 80S ribosome average, EMD-15469; neuronal MT average, EMD-15470. Proteomics data has been submitted to PRIDE with accession number PXD035700. Tomogram preprocessing (https://github.com/williamnwan/TOMOMAN) and PySeg scripts (https://github.com/anmartinezs/pyseg_system) are available on GitHub.

## References

1. M. Bentley, G. Banker, The cellular mechanisms that maintain neuronal polarity. Nat. Rev. Neurosci. 17, 611–622 (2016).

2. N. Hirokawa, S. Niwa, Y. Tanaka, Molecular Motors in Neurons: Transport Mechanisms and Roles in Brain Function, Development, and Disease. Neuron 68, 610–638 (2010).

3. A. Akhmanova, M. O. Steinmetz, Tracking the ends: A dynamic protein network controls the fate of microtubule tips. Nat. Rev. Mol. Cell Biol. 9, 309–322 (2008).

4. J. A. Cooper, Cell biology in neuroscience: mechanisms of cell migration in the nervous system. J. Cell Biol. 202, 725–34 (2013).

5. J. Jaworski, et al., Dynamic Microtubules Regulate Dendritic Spine Morphology and Synaptic Plasticity. Neuron 61, 85–100 (2009).

6. F. Larti, et al., A defect in the CLIP1 gene (CLIP-170) can cause autosomal recessive intellectual disability. Eur. J. Hum. Genet. 23, 331–6 (2015).

7. J. C. Bulinski, Microtubules and Neurodegeneration: The Tubulin Code Sets the Rules of the Road. Curr. Biol. 29, R28–R30 (2019).

8. A. K. Srivastava, C. E. Schwartz, Intellectual disability and autism spectrum disorders: Causal genes and molecular mechanisms. Neurosci. Biobehav. Rev. 46, 161–174 (2014).

9. S. Bodakuntla, A. S. Jijumon, C. Villablanca, C. Gonzalez-Billault, C. Janke, Microtubule-Associated Proteins: Structuring the Cytoskeleton. Trends Cell Biol. 29, 804–819 (2019).

10. S. W. Manka, C. A. Moores, Microtubule structure by cryo-EM: Snapshots of dynamic instability. Essays Biochem. 62, 737–751 (2018).

11. R. B. Dye, S. P. Fink, R. C. Williams, Taxol-induced flexibility of microtubules and its reversal by MAP-2 and Tau. J. Biol. Chem. 268, 6847–6850 (1993).

12. H. Felgner, et al., Domains of neuronal microtubule-associated proteins and flexural rigidity of microtubules. J. Cell Biol. 138, 1067–1075 (1997).

13. J. M. Bassot, R. Martoja, Données histologiques et ultrastructurales sur les microtubules cytoplasmiques du canal éjaculateur des insectes orthoptères. Zeitschrift für Zellforsch. und Mikroskopische Anat. 74, 145–181 (1966).

14. B. A. Afzelius, Microtubules in the spermatids of stick insects. J. Ultrastruct. Res. Mol. Struct. Res. 98, 94–102 (1988).

15. O. Behnke, Incomplete microtubules observed in mammalian blood platelets during microtubule polymerization. J. Cell Biol. 34, 697–701 (1967).

16. C. Bouchet-Marquis, et al., Visualization of cell microtubules in their native state. Biol. Cell 99, 45–53 (2007).

17. B. K. Garvalov, et al., Luminal particles within cellular microtubules. J. Cell Biol. 174, 759–765 (2006).

18. S. Chakraborty, M. Jasnin, W. Baumeister, Three-dimensional organization of the cytoskeleton: A cryo-electron tomography perspective. Protein Sci. 29, 1302– 1320 (2020).

19. N. Schrod, et al., Pleomorphic linkers as ubiquitous structural organizers of vesicles in axons. PLoS One 13, e0197886 (2018).

20. P. C. Hoffmann, et al., Electron cryo-tomography reveals the subcellular architecture of growing axons in human brain organoids. Elife 10:e70269 (2021).

21. H. E. Foster, C. Ventura Santos, A. P. Carter, A cryo-ET survey of microtubules and intracellular compartments in mammalian axons. J. Cell Biol. 221, e202103154 (2022).

22. J. Atherton, M. Stouffer, F. Francis, C. a. Moores, Visualising the cytoskeletal machinery in neuronal growth cones using cryo-electron tomography. J. Cell Sci. 135, jcs259234 (2022).

23. P. R. Burton, Luminal material in microtubules of frog olfactory axons: structure and distribution. J. Cell Biol. 99, 520–528 (1984).

24. E. L. R. Echandía, R. S. Piezzi, E. M. Rodríguez, Dense-core microtubules in neurons and gliocytes of the toad Bufo arenarum Hensel. Am. J. Anat. 122, 157– 167 (1968).

25. J. Atherton, M. Stouffer, F. Francis, C. A. Moores, Microtubule architecture in vitro and in cells revealed by cryo-electron tomography. Acta Crystallogr. Sect. D Struct. Biol. 74, 572–584 (2018).

26. S. Chakraborty, J. Mahamid, W. Baumeister, Cryoelectron Tomography Reveals Nanoscale Organization of the Cytoskeleton and Its Relation to Microtubule Curvature Inside Cells. Structure 28, 991–1003.e4 (2020).

27. F. J. McNally, A. Roll-Mecak, Microtubule-severing enzymes: From cellular functions to molecular mechanism. J. Cell Biol. 217, 4057–4069 (2018).

28. D. Odde, Diffusion inside microtubules. Eur. Biophys. J. 27, 514–520 (1998).

29. P. Guedes-Dias, E. L. F. Holzbaur, Axonal transport: Driving synaptic function. Science 366, eaaw9997 (2019).

30. X. Wang, et al., Cryo-EM structure of cortical microtubules from human parasite Toxoplasma gondii identifies their microtubule inner proteins. Nat. Commun. 12, 3065 (2021).

31. M. Ichikawa, et al., Tubulin lattice in cilia is in a stressed form regulated by microtubule inner proteins. Proc. Natl. Acad. Sci. U. S. A. 116, 19930–19938 (2019).

32. M. Owa, et al., Inner lumen proteins stabilize doublet microtubules in cilia and flagella. Nat. Commun. 10, 1143 (2019).

33. D. Zabeo, et al., A lumenal interrupted helix in human sperm tail microtubules. Sci. Rep. 8, 2727 (2018).

34. M. R. Leung, et al., Structural specializations of the sperm tail. Cell 186, 2880–2896.e17 (2023).

35. M. LeDizet, G. Piperno, Identification of an acetylation site of Chlamydomonas alpha-tubulin. Proc. Natl. Acad. Sci. U. S. A. 84, 5720–5724 (1987).

36. Z. Xu, et al., Microtubules acquire resistance from mechanical breakage through intralumenal acetylation. Science 356, 328–332 (2017).

37. D. Portran, L. Schaedel, Z. Xu, M. Théry, M. V. Nachury, Tubulin acetylation protects long-lived microtubules against mechanical ageing. Nat. Cell Biol. 19, 391–398 (2017).

38. E. A. Katrukha, D. Jurriens, D. M. Salas Pastene, L. C. Kapitein, Quantitative mapping of dense microtubule arrays in mammalian neurons. Elife 10:e67925 (2021).

39. I. Topalidou, et al., Genetically separable functions of the MEC-17 tubulin acetyltransferase affect microtubule organization. Curr. Biol. 22, 1057–1065 (2012).

40. A. Matsuyama, et al., In vivo destabilization of dynamic microtubules by HDAC6-mediated deacetylation. EMBO J. 21, 6820–6831 (2002).

41. Y. Zilberman, et al., Regulation of microtubule dynamics by inhibition of the tubulin deacetylase HDAC6. J. Cell Sci. 122, 3531–3541 (2009).

42. C. Cuveillier, et al., MAP6 is an intraluminal protein that induces neuronal microtubules to coil. Sci. Adv. 6, eaaz4344 (2020).

43. H. Sosa, D. Chrétien, Relationship between Moire patterns, tubulin shape, and microtubule polarity. Cell Motil. Cytoskeleton 40, 38–43 (1998).

44. L. G. Tilney, et al., Microtubules: Evidence for 13 protofilaments. J. Cell Biol. 59, 267–275 (1973).

45. P. W. Baas, J. S. Deitch, M. M. Black, G. A. Banker, Polarity orientation of microtubules in hippocampal neurons: uniformity in the axon and nonuniformity in the dendrite. Proc. Natl. Acad. Sci. 85, 8335–8339 (1988).

46. A. Martinez-Sanchez, et al., Template-free detection and classification of membrane-bound complexes in cryo-electron tomograms. Nat. Methods 17, 209– 216 (2020).

47. S. H. W. Scheres, RELION: Implementation of a Bayesian approach to cryo-EM structure determination. J. Struct. Biol. 180, 519–530 (2012).

48. K. Johnson, J. Wall, Structure and molecular weight of the dynein ATPase. J. Cell Biol. 96, 669–678 (1983).

49. L. A. Amos, J. Löwe, How Taxol stabilises microtubule structure. Chem. Biol. 6, R65–9 (1999).

50. I. Poser, et al., BAC TransgeneOmics: A high-throughput method for exploration of protein function in mammals. Nat. Methods 5, 409–415 (2008).

51. J. Jumper, et al., Highly accurate protein structure prediction with AlphaFold. Nature 596, 583–589 (2021).

52. G. Piperno, M. LeDizet, X. J. Chang, Microtubules containing acetylated alpha-tubulin in mammalian cells in culture. J. Cell Biol. 104, 289–302 (1987).

53. S. Nithianantham, et al., Tubulin cofactors and Arl2 are cage-like chaperones that regulate the soluble αβ-tubulin pool for microtubule dynamics. Elife 4:e08811 (2015).

54. G. Tian, et al., Pathway leading to correctly folded β-tubulin. Cell 86, 287–296 (1996).

55. A. Cook, et al., The structure of the nuclear export receptor Cse1 in its cytosolic state reveals a closed conformation incompatible with cargo binding. Mol. Cell 18, 355–367 (2005).

56. G. Tian, S. Thomas, N. J. Cowan, Effect of TBCD and its regulatory interactor Arl2 on tubulin and microtubule integrity. Cytoskeleton 67, 706–714 (2010).

57. J. W. Francis, L. E. Newman, L. A. Cunningham, R. A. Kahn, A trimer consisting of the Tubulin-specific Chaperone D (TBCD), regulatory GTPase ARL2, and β-tubulin is required for maintaining the microtubule network. J. Biol. Chem. 292, 4336–4349 (2017).

58. M. Serna, et al., The structure of the complex between α-tubulin, TBCE and TBCB reveals a tubulin dimer dissociation mechanism. J. Cell Sci. 128, 1824–1834 (2015).

59. M. McBurney, et al., Differentiation and maturation of embryonal carcinoma-derived neurons in cell culture. J. Neurosci. 8, 1063–1073 (1988).

60. M. M. Falconer, U. Vielkind, D. L. Brown, Association of acetylated microtubules, vimentin intermediate filaments, and MAP 2 during early neural differentiation in EC cell culture. Biochem. Cell Biol. 67, 537–544 (1989).

61. C. Coombes, et al., Mechanism of microtubule lumen entry for the α-tubulin acetyltransferase enzyme αTAT1. Proc. Natl. Acad. Sci. U. S. A. 113, E7176– E7184 (2016).

62. D. Chretien, F. Metoz, F. Verde, E. Karsenti, R. H. Wade, Lattice defects in microtubules: Protofilament numbers vary within individual microtubules. J. Cell Biol. 117, 1031–1040 (1992).

63. M. Cyrklaff, et al., Cryoelectron tomography reveals periodic material at the inner side of subpellicular microtubules in apicomplexan parasites. J. Exp. Med. 204, 1281–1287 (2007).

64. S. Reber, A. A. Hyman, Emergent Properties of the Metaphase Spindle. Cold Spring Harb. Perspect. Biol. 7, a015784 (2015).

65. M. Javier-Torrent, G. Zimmer-Bensch, L. Nguyen, Mechanical Forces Orchestrate Brain Development. Trends Neurosci. 44, 110–121 (2021).

66. E. W. Dent, S. L. Gupton, F. B. Gertler, The growth cone cytoskeleton in Axon outgrowth and guidance. Cold Spring Harb. Perspect. Biol. 3, 1–39 (2011).

67. K. J. Verhey, J. Gaertig, The Tubulin Code. Cell Cycle 6, 2152–2160 (2007).

68. S. A. Lewis, G. Tian, N. J. Cowan, The [alpha]- and [beta]-tubulin folding pathways. Trends Cell Biol. 7, 479–484 (1997).

69. L. M. Pinho-Correia, A. Prokop, Maintaining essential microtubule bundles in meter-long axons: a role for local tubulin biogenesis? Brain Res. Bull. 193, 131– 145 (2023).

70. R. J. Lasek, C. Dabrowski, R. Nordlander, Analysis of axoplasmic RNA from invertebrate giant axons. Nat. New Biol. 244, 162–165 (1973).

71. R. B. Campenot, K. Lund, D. L. Senger, Delivery of newly synthesized tubulin to rapidly growing distal axons of rat sympathetic neurons in compartmented cultures. J. Cell Biol. 135, 701–709 (1996).

72. A. Bhamidipati, S. a. Lewis, N. J. Cowan, ADP ribosylation factor-like protein 2 (Arl2) regulates the interaction of tubulin-folding cofactor D with native tubulin. J. Cell Biol. 149, 1087–1096 (2000).

73. I. Gasic, T. J. Mitchison, Autoregulation and repair in microtubule homeostasis. Curr. Opin. Cell Biol. 56, 80–87 (2019).

74. M. Srayko, E. T. O’Toole, A. A. Hyman, T. Müller-Reichert, Katanin Disrupts the Microtubule Lattice and Increases Polymer Number in C. elegans Meiosis. Curr. Biol. 16, 1944–1949 (2006).

75. L. J. Davis, D. J. Odde, S. M. Block, S. P. Gross, The importance of lattice defects in katanin-mediated microtubule severing in vitro. Biophys. J. 82, 2916–2927 (2002).

76. A. Aher, et al., CLASP Mediates Microtubule Repair by Restricting Lattice Damage and Regulating Tubulin Incorporation. Curr. Biol. 30, 2175–2183.e6 (2020).

77. W. Wang, et al., Gigaxonin interacts with tubulin folding cofactor B and controls its degradation through the ubiquitin-proteasome pathway. Curr. Biol. 15, 2050–2055 (2005).

78. R. Parvari, et al., Mutation of TBCE causes hypoparathyroidism-retardation-dysmorphism and autosomal recessive Kenny-Caffey Syndrome. Nat. Genet. 32, 448–452 (2002).

79. E. Flex, et al., Biallelic Mutations in TBCD, Encoding the Tubulin Folding Cofactor D, Perturb Microtubule Dynamics and Cause Early-Onset Encephalopathy. Am. J. Hum. Genet. 99, 962–973 (2016).

80. S. Edvardson, et al., Infantile neurodegenerative disorder associated with mutations in TBCD, an essential gene in the tubulin heterodimer assembly pathway. Hum. Mol. Genet. 25, 4635–4638 (2016).

81. D. N. Mastronarde, Automated electron microscope tomography using robust prediction of specimen movements. J. Struct. Biol. 152, 36–51 (2005).

82. W. J. H. Hagen, W. Wan, J. A. G. Briggs, Implementation of a cryo-electron tomography tilt-scheme optimized for high resolution subtomogram averaging. J. Struct. Biol. 197, 191–198 (2017).

83. J. Zivanov, et al., New tools for automated high-resolution cryo-EM structure determination in RELION-3. Elife 7:e42166 (2018).

