## Supplementary Data for "Cryo-electron tomography suggests tubulin chaperones form a subset of microtubule lumenal particles with a role in maintaining neuronal microtubules"

**This PDF file includes:**

SI Methods
Figures S1 to S11
Tables S1 to S2
Legends for Movies S1 to S5
SI References

**Other supporting materials for this manuscript include the following:**

Movies S1 to S5

**SI Methods**:

**Primary neuron culture and grid preparation**

Rat hippocampal neurons derived from embryonic rats (E17-21) in accordance with the procedures accepted by the Max Planck Institute for Biochemistry were kindly provided by Baumeister group members (1). Briefly, the embryo tissue was dissected and minced in HBSS-HEPES buffer and trypsinized for 15-30 mins to collect hippocampal neurons as described elsewhere (1). Collected neurons were washed with DMEM and 10% FBS (Invitrogen), and seeded on 4-well 35 mm dishes to a concentration of $5×10^5$ cells/well each containing four poly-L-lysine coated R1/4 Au 200 mesh Quantifoil grids (Quantifoil Micro Tools). Cells were kept in an incubator at 37 °C and 5% $CO_2$ for ~4 h. Then the grids were transferred to a culture dish containing pre-conditioned B27 medium. After 3 days from seeding, cytosine β-D-arabinofuranoside (Ara-C) (SIGMA, C1768) was added (final concentration 5 µM) to the culture dish to stop glial cell duplication. Neurons were cultivated for 10–18 days *in vitro* (DIV). One third of the culture medium was exchanged once a week. Grids were vitrified on day 12 using a Vitrobot Mark 4. The Vitrobot was set to 37°C, 90 % humidity, blot force 10, blot time of 10 s and 2 s drain time.

**P19 cell culture, neuronal differentiation and grid preparation**

Murine pluripotent P19 cells (DSMZ, ACC 316) were cultured in P19 growth medium (P19GM) containing alpha-MEM (Gibco, 12571063) supplemented with nucleosides, 10% (v/v) FBS (Thermo Fisher Scientific, 16000036), 100 mg/mL each of penicillin and streptomycin (Gibco, 15140122) at 37°C with 5% $CO_2$. Sample preparation on EM grids was essentially same as described before (2). Briefly, Pluripotent P19 cells were grown on plasma cleaned, UV sterilized Gold Quantifoil grids (R2/1, Au 200 mesh grid) that had an additional 20 nm carbon. Cells were kept in an incubator overnight to allow adhesion, followed by vitrification by plunge freezing using a Vitrobot Mark 4 (Thermo Fisher Scientific). In all cases, Vitrobot was set to 37°C, 95 % humidity, blot force 10, blot time of 10 s and 2 s drain time.

In order to analyze lumenal particle entry to MT lumen, 15 µM Nocodazole (Sigma-Aldrich, M1404) was added to pluripotent P19 cells grown on EM grids in cell culture media for 30 mins. Media containing Nocodazole was deposed and cells were washed with PBS three times to remove residual drug followed by addition of fresh media to induce fresh MT polymerization. Grids were vitrified after 15 mins of incubation.

Pluripotent P19 cells were induced to differentiate into neurons in induction medium (P19IM) containing 95% alpha-MEM and 5% FBS. Briefly, cells were collected after trypsinization with 0.05% trypsin and re-suspended in P19IM. Cells were then allowed to aggregate in a low binding petri-dish in presence of 0.5 µM all-trans retinoic acid (RA) (SIGMA, R2625-50MG). Aggregates of P19 cells formed within 2 days and further continued for another 2 days with fresh P19IM containing RA. Aggregates were then harvested and treated with 0.05% trypsin (Gibco, 25300054) and DNase (50 µg/mL) (New England Biolabs GmbH, M0303S) followed by resuspension in P19GM. Cells were plated in 35 mm poly-L-lysine-coated tissue culture dishes at a density of $~5×10^5$ cells/ml each containing four poly-L-lysine coated R2/1 Au Quantifoil 200 mesh grids. Ara-C (SIGMA, C1768) was added at 5 mM concentration after 2 days and culture continued for 7-10 days. Grids were vitrified between days 10-14.

**hiPSCs culture, neuronal differentiation and grid preparation**

Human iPSCs (male CESCG-297) were kindly provided by Michael Snyder's laboratory at Stanford University. hiPSCs were derived from peripheral blood mononuclear cells following Institutional Review Board (IRB) approval and Stem Cell Research Oversight (SCRO) approval at Stanford University and under informed consent. hiPSCs were maintained below passage 40, regularly tested for mycoplasma (assayed with Mycoplasma PCR Detection Kit, iNtRON), and examined for genome integrity (using low coverage whole genome sequencing). hiPSCs were cultured in vitronectin (Thermo Fisher Scientific, A14700) coated dish (0.5 µg/cm$^2$) with Essential 8 medium with supplement (Thermo Fisher Scientific, A1517001). For neuronal differentiation, we adapted and modified published methods that generate excitatory neurons using rapid Neurogenin-2 (NGN2) induction (3, 4). Briefly, hiPSCs were transduced with TetO-hNGN2-P2A-eGFP-T2A-PuroR and rtTA lentivirus at 60% of confluency (Day 0). After 24 h, the virus-containing medium was washed away and replaced with 1:1 ratio of Neurobasal A (Thermo Fisher Scientific, 10888022) and DMEM/F12 medium containing GlutaMAX, Insulin (Gibco, 51300-044), N2/B27 without Vitamin A (Thermo Fisher Scientific, 17502048/12587010) supplements, and 1 µg/ml of doxycycline to activate TetO promoter (Day 1). hiPSCs expressing hNGN2-P2A-eGFP-T2A-PuroR were selected by adding 3 µg/ml puromycin (Day 2) for 24 h. Cells were then dissociated with Accutase (Thermo Fisher Scientific, A1110501), counted, and re-plated in the triple coated dish containing EM grids described below (Day 3 - seeding), and maintained in Neurobasal medium (Thermo Fisher Scientific, 21103049) containing B-27 w/o Vitamin A (Gibco, 12587010), GlutaMAX (Thermo Fisher Scientific, 35050061), 200 µM of L-Ascorbic Acid (Sigma, A5960), 1 µM of DAPT (Tocris, 2634), 1 µg/ml puromycin, and 1µg/ml doxycycline. The triple coating is composed of 0.1 mg/ml of Poly-D-Lysine (Sigma, P0899), 10 µg/ml of Laminin (Sigma, 11243217001), and 10 µg/ml of Fibronectin (Sigma, F0895). Cells were cultured for two more days with 1 µg/µl puromycin and 1 µg/ml doxycycline (Day 5). Half of the medium was changed every two days without puromycin and doxycycline until the induced neurons were vitrified (Day 9).

Lentivirus generation: two lentivirus plasmids from a published study (4), each containing hNGN2-P2A-eGFP-T2A-PuroR under the control of TetO promoter (Addgene 79823) and reverse tetracycline activator (rtTA) (Addgene 19780), were used to produce transgenes. Lentivirus was generated by co-transfecting transgene plasmid with two helper plasmids, pMD2 (Addgene 12259) and psPAX2 (Addgene 12260), by calcium phosphate transfection kit (Takara, 631312) following the manufacturer's protocol into HEK293 cells. After 48-72 h of incubation, media containing lentiviruses were harvested and concentrated at 112,000 g for 3 h, aliquoted, and stored at -80°C.

EM Grid micro-patterning: gold (Au) 200-mesh grids with a holey 12 nm thick film made of SiO2 (R1/4, hole size/spacing in µm, Quantifoil Micro Tools), were micro-patterned (5). Briefly, grids were plasma cleaned on both sides at 100 W power with a 10 cm$^3$ per min flow rate of oxygen gas for

30–40 s (Diener Femto Plasma, DienerElectronic). Next, grids were incubated on droplets of poly-l-lysine grafted with polyethylene glycol (PLL(20)- g[3.5]-PEG(5), SuSoS AG) at a concentration of 0.5 mg/ml in 10 mM HEPES pH 7.4, for 1h at room temperature on parafilm in a humid chamber. Following passivation, the grids were blotted with filter paper from the back, allowed to dry for few seconds and immediately placed on a drop of PLPP (4-benzoylbenzyl-trimethylammonium chloride, 14.5 mg/ml) in a sealed glass bottom ibidi µ-Dish 35mm low (ibidi). Grids were immediately micro-patterned using a DMD-based illumination (Primo, Alvéole Lab), applying a dose of 1,500 $mJ/mm^2$. Grids were promptly retrieved from the PLPP solution, washed in a 1,000 µl drop of water and two consecutive washes in 300 µl drops of PBS supplemented with Ca and Mg. Patterned grids were stored wet in PBS supplemented with Ca and Mg at 4 °C in a humid chamber. For functionalization of the micropatterns, grids were incubated at room temperature for 30 min in a 20 µl drop of 50 µg/ml laminin (Sigma, 11243217001) and 50 µg/ml fibronectin (Thermo Fisher Scientific) in PBS supplemented with Ca and Mg. Grids were then washed three times in 300 µl drops of the corresponding buffer and stored at 4 °C in a humid chamber until use. hiPSC cells were seeded on laminin/fibronectin micro-patterned grids at a density of $2\times10^4$ cells per $cm^2$ in a glass bottom ibidi µ-Dish 35mm high (ibidi), and incubated for 48 h in this dish. In parallel, another glass bottom ibidi µ-Dish 35mm high (ibidi) was seeded with $2\times10^5$ cells per $cm^2$ (without grids). After 48 h, the grids were transferred (from the $2\times10^4$ cells per $cm^2$ seeded dish) to the $2\times10^5$ cells per $cm^2$ seeded dish. Neurons were grown for 6 days post-seeding with half-media change every 2 days. Grids were plunged on day 6 post-seeding using a Leica GP plunger at a 70% humidity and blotted for 1.5 sec, no nanogold fiducials were added.

Micropattern design: micropatterns were designed in Inkscape (http://www.inkscape.org/) as 8-bit binary files and exported as .png files, which can be loaded into the Leonardo software v.4.12 (Alvéole Lab). Neuronal circuits micropatterns used for hiPSCs were designed such that 10 µm circles were patterned at the center of the grid square to direct cell body attachment. Circles were connected alternatingly with straight or curly lines (10 µm wide) to facilitate extension of neuronal process. Micro-patterned circuit covered an area of 10 x 8 grid squares (in a 200-mesh grid).

**HeLa cell culture, grid preparation and FIB micromachining**

HeLa Kyoto cells expressing both green fluorescent protein (GFP)-tagged β-tubulin and mCherry tagged histone (H2B-mCherry) were kindly provided by Anthony Hyman's laboratory at MPI-CBG, Dresden. Cells were cultured at 37°C with 5% $CO_2$ in Dulbecco's modified Eagle's medium (DMEM) (Gibco, 31966021) containing 10% (v/v) fetal bovine serum (FBS) (Gibco, 16000036), 2 mM L-glutamine, 100 mg/mL penicillin, 100 mg/mL streptomycin, and 0.5 mg/mL geneticin (G418) (Gibco, 10131019).

For the preparation of mitotic HeLa cells, mitotically synchronized cells using thymidine (Sigma, T1895-5G) block as described before (2) were detached from the culture flask by mechanical shake off, and the collected medium was centrifuged at 300 g for 2 min to concentrate the cell pellet. 4 µL of the cell suspension was deposited onto plasma cleaned holey carbon-coated 200 mesh copper R 2/1 grids followed by vitrification by plunge freezing using a Vitrobot Mark 4 (Thermo Fisher Scientific) using similar conditions as in P19 sample preparation.

Plunge-frozen grids fixed into custom-made autogrids were mounted into a 45°-pre-tilted shuttle and micro-machined using a dual-beam (FIB/SEM) microscope (Quanta 3D FEG, FEI) as described before (2, 6). For improved conductivity of the final lamella, grids were sputter coated after cryo-FIB preparation with platinum in the Quorum prep-chamber (10 mA, 3 s).

**Cryo-TEM and tomography**

Tilt series were collected on a Titan Krios (Thermo Fisher Scientific) operated at 300 kV equipped with a field-emission gun, a Quantum post-column energy filter (Gatan) operated in the zero-loss mode using a 20 eV slit, a K2 Summit direct detector camera (Gatan) operated in counting mode, and/or a Volta phase plate (VPP) (Thermo Fisher Scientific) for part of the data. Alignment and operation of the Volta phase plate were essentially carried out as described by Fukuda et al. (7), applying a beam tilt of 10 mrad for autofocusing. Data were recorded with SerialEM software (8). Only the best quality tomograms with good contrast produced from cellular areas thinner than 200 nm were used for image processing (**Table S1**).

Tilt series of primary neurons were acquired either with dose-symmetric (9) or asymmetric bi-directional tilt scheme and a nominal defocus ranging from 2 to 6 µm at an EFTEM magnification of ×64,000 resulting in pixel size at the specimen level of 2.24 Å. Total dose was kept under 100 e/Å$^2$. 60 tomograms were processed for subsequent structural analysis.

Tilt series of hiPSC-derived neurons were collected with dose symmetric scheme at an EFTEM magnification of ×64,000 corresponding to pixel size at the specimen level of 2.129 Å, 2.5–4µm defocus, with constant dose for all tilts, total dose under 150 e/Å$^2$. Tilt-series were acquired with the VPP. 8 tomograms were processed for subsequent structural analysis.

A total of 46 bi-directional tilt-series corresponding to pluripotent control, Taxol-treated and Nocodazole-treated P19 cells was collected at ×42,000 resulting in pixel size of 3.42 Å using VPP with a target defocus of 0-200 nm. A total of 35 dose-symmetric tilt series corresponding to pluripotent and differentiated P19 cells was also collected with an EFTEM magnification of ×64,000 corresponding to pixel size at the specimen level of 2.24 Å and 2-5 µm defocus. In all cases, cumulative dose ranged between 90-120 e/Å$^2$. HeLa cell tomograms were collected from FIB-lamellae at ×33,000 resulting in pixel size of 4.21 Å using VPP with a target defocus of 0 nm as described previously (2).

**Processing of tilt-series**

For preprocessing of tilt series, the MATLAB based TOMOgram MANager (TOMOMAN) package was used (10). Frames were first sorted, followed by motion correction using MontionCor2 (11) with 3 × 3 patches. Bad frames were then omitted from the motion corrected stacks. Frames were exposure filtered according to their cumulative electron dose. Defocus parameters were determined using GCTF 1.06 (12) or CTFFIND 4.1.14 (13) on unfiltered stacks. For high resolution data processing, 3D defocus gradients in primary neuron and P19 tomograms were corrected using NovaCTF (14). CTF correction via phase-flipping used a Z-slab thickness of 15 nm. The resulting tomograms were consecutively binned via Fourier cropping. Preprocessing of hiPSC data was performed using Warp 1.7.b (15) including frame alignment, gain correction, dose filtering, tilt sorting, and CTF fitting. In-focus tomograms collected with VPP were not CTF corrected. In all cases, the dose-filtered stacks were aligned with patch tracking in the IMOD software package (16). Final alignment of the tilt-series images was performed using the linear interpolation option in IMOD without CTF correction. For tomographic reconstruction by back-projection, the radial filter options were left at their default values (cut off, 0.35; fall off, 0.05). Missing wedge compensated and denoised cross-section of the tomogram represented in **Fig. S1A** and **S1D** was prepared for visualization purpose using IsoNet (17).

**MT tracing and curvature measurement**

MT segmentations were performed with the automated module in Amira software v.6.2.0 (Thermo Fisher Scientific) as described before (2) in the IMOD reconstructed tomograms using a generic missing wedge-modulated hollow cylinder with outer and inner radii of 14 and 7 nm, respectively and 120 nm length as a template. The coordinates of the traced filaments were resampled in MATLAB (MathWorks) to obtain equidistant points. MT curvature was determined by measuring the tangent-correlation length as described in our previous work (2). Data analysis was done by fitting a linear regression line in Origin 9.2 (OriginLab, Northampton, MA) on the data points obtained from each tomogram. ${}^{a}L_{p}$ values were calculated from the inverse slopes of the linear regression lines. Goodness of fits were determined from the Adjusted-$R^2$ values. Only fits with Adj-$R^2$>0.8 are considered.

**Density tracing and particle picking**

For density tracing and particle picking, tomograms were smoothed by Gaussian low-pass filtering at σv=0.5 pixels and MT lumen is segmented based on the MT centerline tracing. Densities within the lumen were traced in 3D using a discrete Morse theory based software package PySeg (18). Briefly, grayscale minima positions were identified and the most relevant ones were preserved by topological persistence simplification, while discarding those generated from noise or spurious densities. Next, mean shift clustering method (19) with 6 nm bandwidth was applied that allowed detection of particles and particle picking within MT lumen in an unbiased template-free manner. The persistence threshold was set for picking all lumenal particles independent of the type of density. Therefore, this threshold was adjusted for having the same density of minima at the shell for every MT, specifically 0.004 vertices/nm3 for phase plate tomograms and 0.006 vertices/nm3 for the defocus tomograms.

**Subtomogram averaging of lumenal particles**

MTs were segmented using the filament-tracing module in Amira v.6.2.0, (Thermo Fisher Scientific), and lumenal particles picked as described above. Subtomogram analysis was performed using RELION (v.3.0.5) following published protocols (20). During the refinement, particle half-datasets were processed independently. Unless stated explicitly, all refinements were performed *de novo*, that is, without the use of external references. Datasets corresponding to different cells were processed separately (**Table S1**).

Lumenal particle sub-tomograms from the coordinates determined by the automated particle picking algorithm were extracted at 1x binning (2.24 Å/pix for all neuronal samples, except 2.129 Å/pix in case of hiPSC data) within a box size of 128 pixels$^3$. For VPP data corresponding to P19 and HeLa, no CTF correction was performed and lumenal particle sub-tomograms were reconstructed using IMOD (3.42 Å/pix and 4.21 Å/pix) within a box size of 128 pixels$^3$. For P19 cell data, we combined VPP datasets from different conditions to increase the particle number and improve angular sampling. All the subtomograms correspond to side views only on the MTs and lacked views from cross-sections. Nevertheless, individual particles having different orientation within the MT lumen helped mitigate this problem.

Here, the STA procedure for neuronal samples is described. All other samples were analyzed using the same strategy. 50,910 subtomograms were first combined with randomized Euler angle resulting in a sphere (**Fig. S4**, top left) and auto-refined in RELION with global search to shift them to a common origin (**Fig. S4**, top right) using a soft cylindrical mask. This initial model was used as an input for 3D classification with a global spherical mask (130 Å diameter). In order to resolve heterogeneity in the dataset, a multi-tier (at least three) and exhaustive classification strategy was used. For the first classification step, particle alignment was skipped and large number of classes (~20) were used to pick up small differences. Multiple different classification runs were performed using different RELION optimization parameters (class numbers 5–25, T values 1–8, limit resolution E-step 8–12 Å) for comparison. Based on the classification, subtomograms were sorted and 3D auto-refined using global search parameters to generate class averages. In the second classification step, refined class averages from the first classification run were used as input (**Fig. S4)** with similar classification settings as in the previous run. We followed this procedure sequentially until no new classes were obtained from the refined class averages of the previous run. The classes that did not produce any new structure were taken as the final 'pure' class averages. Similarity between the class averages was apparent (containing a ring-like density), although differences were observed in the orientation and position of the small globular densities on the ring. Therefore, the class averages were not mixed and treated as separate classes.

For further validation of the classification procedure, the entire neuronal dataset was divided into 6 subsets based on the acquisition date, and processed individually as described above. All reported classes were retrieved from these subsets.

The small size of the complexes and low resolution could introduce starting model bias, as well as produce non-optimum refinement solutions. Therefore, subsequent validation runs were performed for each 'pure' class average to check whether similar structures are obtained when a soft-edged sphere or a cylinder were used as starting model instead of the corresponding particle average (**Fig. S3, S4B-D**). If sorted subtomograms converged to a similar structure, the class average was taken to represent an optimum-refinement solution. In order to assess similarity between classes, we calculated cross-correlations between the final solutions obtained from the particle average as starting model and those of obtained from simple sphere and/or cylinder in Chimera (v1.15) (**Fig. S4B-D**) (21). Cross-correlations were obtained using the 'fit' command in the command line and further refined using 'fit in map' from the menu.

Finally, the class averages were locally refined using a soft particle masks and post processed to determine gold standard FSC curve.

**Particle concentration measurements**

Overall lumenal particle concentrations present in different neuronal samples were measured by counting traced particles obtained by PySeg within each MT of the corresponding samples. Concentration is measured as number of particles per $\mu m^3$ of lumenal volume. Prior to counting all particle, coordinates were refined in RELION. Following our rigorous 3D-classification method described above, concentrations of the individual class averages in their respective samples were determined by mapping them back into their MT of origin followed by counting their occurrences in each MT across tomograms belonging to particular cell type or conditions and converting them into mean number of particles/ $\mu m^3$ of lumenal volume. The whole process was done semi-automatically. In all cases, lumen radius was set to 6 nm and MT length was obtained from the segmentations. MTs with length less than 100 nm was excluded from the analysis.

**Subtomogram averaging and polarity determination of MTs**

Resampled coordinates of the primary neuronal MTs extracted from Amira were used to make a tubular grid in MATLAB by placing points on each tubulin subunit taking into account inter-subunit distance of 40 Å along the filament z-axis and 60 Å around the helix. These positions were oversampled 2x-times and subtomograms were extracted from these grid points. Subtomograms extracted from the same filament were assigned a class number which helped in determination of MT polarity. Subtomogram averaging was performed in STOPGAP (22). An initial reference was generated *de novo* from a single MT filament. 10,076 subtomograms were extracted from the oversampled positions using a binning factor of 2 (8.96 Å/pixel) and a box size of 64 pixels$^3$ and a starting reference was generated by averaging all sub-tomograms. Due to in-plane randomization, the initial structure resembles a featureless cylinder. Several rounds of alignment of the in-plane angle were conducted. Shift refinements were limited to 4 and 6 nm in x- and y- direction along the filament centerline. This averaging approach resulted in the emergence of the helical pattern of the MT (inset: **Fig. S1B, E**). We used this optimized starting model for refining the whole data-set for 5-6 iterations sequentially at bin4, bin2, and bin1. In order to determine the polarity of the MTs, each MT in the dataset was averaged separately using multiclass averaging and subtomogram orientations were visualized in Chimera as lattice maps using 'place object' plugin (23). Polarities were sorted out using visual inspection of the pF skew and followed by 180° change of the in-plane angles in the motive list. Finally, 34906 bin1 subtomograms from 19 MTs (in two tomograms) were extracted using a box size of $96^3$ pixels$^3$ and aligned iteratively, resulting in a refined map of a MT at a resolution of 8.19 Å. For fitting the tubulin dimer (PDB**:** 1tub**)** into the obtained map, we used Chimera for rigid body fitting however, fitting was not optimal. Therefore, we used qwikMD (24) with default parameters for flexible fitting of the tubulin dimer that is represented in **Fig.1H**.

**Mass measurement of lumenal particles**

The luminal particles with a voxel size of 2.24/3.42 Å were filtered using a circular kernel (TomToolbox, tom_filter kernel=6) (25). The threshold for binarization was determined by measuring the mass of the ribosome particles from the same tomogram as described in literature (26). The threshold was decreased until the mean mass of all ribosome particles was ~3.6 MDa,(27) and then applied to the luminal particles for binarization. To correct for the different contrast regimes a fudge factor of 0.5 was applied to the threshold. Binarized luminal particles were segmented using Matlab's regionprops3, yielding the number of voxels for the biggest object inside one subvolume. The corresponding mass of each particle was calculated based on voxel size and the protein density (1.3 g $cm^{-3}$).

**Template matching and subtomogram averaging of ribosomes**

To generate a subtomogram average of the ribosomes present in the neurons and P19 cells, ribosomes were localized by template matching in PyTom (28). As a template, we used the 4 nm down filtered structures of 80S ribosome (EMD: 1480) rescaled to the tomogram pixel size. False positives were identified by constrained cross-correlation (CCC) histogram which showed well separated peaks for the true and false positives (**Fig. S5C**). However, picked positions were also manually verified. In case of P19 ribosomes, 5553 (at bin-1, 3.42 Å pixel size) subtomograms were extracted at the detected coordinates with their third Euler angle randomized and refined in RELION, which converged into a ribosome density (**Fig. S5D**). Particles were next sorted in 3D classification steps in RELION to remove remaining false positives. After final refinement and post processing, we obtained a density map at 18 Å resolution (**Fig. S5E**) from 4723 particles. In case of neuronal ribosomes, 975 4x binned subtomograms centered on the identified coordinates were extracted and refined in RELION using a data-driven template. The refinement converged into a ribosome density. The data was further classified to remove false positives. All the good classes were combined and un-binned sequentially to 2x and 1x and final refinement gave 28 Å resolution ribosome average.

**Polyribosome detection**

Polyribosome detection was essentially based on hierarchical transformation clustering as described recently in Jiang et al. (29). The algorithm considered the directions of the transformations (rotations and translations). The method consists of 3 major steps: (i) transformations (rotations, translations) between ribosome pairs were calculated; (ii) distance matrices for rotations and translations were computed and combined; (iii) transformations were clustered based on the combined distance matrix (**Fig. S5G-I**).

**Heat Maps of lumenal particle micro-clustering**

Heat Maps (**Fig**. **7D**, **7I**) were calculated by generating a volume of zeros with the size of the analyzed tomograms (928x928x500 voxels). For each particle position, the voxel value in the empty volume was set to 1. The 3D heat map was calculated by moving a spherical kernel with 2x radius of the particle over the volume. The mean for each kernel position was determined. Finally the 3D heat map was projected in z.

**Spatial statistics of lumenal particle organization**

We computed 1$^{st}$ order univariate (NN-distance) and 2$^{nd}$ order bivariate (Ripley's L) functions for spatial statistical analysis of 3D organization of lumenal particles in a cellular compartment (MT lumen) as proposed in (18). The detailed description of the numerical corrections needed for successfully computation of these functions from cryo-ET data is described in Martinez-Sanchez et al (30). Protein localizations, angular orientations and 3D shapes were taken from STA, MTs lumen were segmented from the traced MT centerlines and a assuming a radius section of 6 nm. Complete Spatial Randomness with Volume exclusion synthetic patterns (CSRV) were used as null-models. In all cases, we simulated 20 instances for each MT to construct the interval of confidences for null-models. One sided Welch tests were used to discard the next null-hypothesis:

experimental particle density has the same mean as the CSRV case at a specific distance to a reference point (MT plus end or breakpoint).

**Statistical analysis**

Statistical analyses were performed with Origin 9.2. All boxes in the box-plot representations are bound by the 25th–75th percentile, whiskers span 5th to 95th percentile. Mean values were marked by a black box. Box plots are embedded inside a violin which indicate the distribution of the data. For statistical significance, nonparametric Mann-Whitney tests were performed unless stated otherwise. Significances were indicated by stars where ** indicated $P<0.01$.

**Visualization and Rigid body fitting**

RELION post-processing were used to calculate FSC and perform B factor sharpening. UCSF Chimera (21) and ChimeraX (31) was used for visualization and figure generation. Numerical data and statistics were analyzed in Origin software. Quantification and plots were done in MATLAB (R2015b, The Mathworks, Inc. Massachusetts, USA). Particle picking and density tracing were visualized using Paraview. Molecular model fitting was performed in Chimera (21). For rigid body fitting, yeast Cse1p (PDB:1Z3H) (32) filtered to 25 Å was used to represent the TBCD as described in (33). Models were fit into non-segmented density maps using the fit command from the command line in an unbiased manner. Since TBCD model is the largest and has distinct shape, it was always fitted first and rests were fitted sequentially. The 30 most probable fits were obtained. Highest cross-correlation hits were selected and refined using 'fit in map' command from the menu.

**Affinity purification and Mass spectrometry (AP-MS) based proteomics**

HeLa Kyoto cells expressing GFP-tagged β-tubulin proteins from BAC transgenes (34) were grown to near-confluency on two 15-cm cell culture dishes per interaction experiment. DSS (Thermo Fisher Scientific, 21555) solution was prepared fresh in dry DMSO at a concentration of 20 mM. Before crosslinking, cells were washed with ice-cold PBS (pH 8.0) to remove amine-containing culture media and extracellular proteins. DSS solution was added at a final concentration of 5 mM for 30 mins. The reaction was quenched by 10 mM Tris-Cl for 15 mins at room temperature. Cells were collected by scraping and snap frozen in liquid $N_2$. Three replicates were harvested. Cell pellets were lysed with a buffer containing 0.1% NP-40 (Sigma) and subjected to affinity purification using anti-GFP antibodies immobilized on a magnetic beads (Miltenyi Biotec, 130-091-125). Purified proteins were digested sequentially with LysC and trypsin, desalted and subjected to mass spectrometric analysis on an Orbitrap instrument (Q Exactive HF) at the mass spectrometry facility of Max-Planck institute of biochemistry. Raw files were processed with MaxQuant (35). MaxQuant output data was processed in Perseus (v.1.6.14.0) program with standard procedure (36). Protein identifications were filtered, removing hits to the reverse decoy database as well as proteins only identified by modified peptides. Protein LFQ intensities were logarithmized and missing values imputed by values simulating noise around the detection limit.

**Protein structure and disorder prediction**

Secondary structure of the AP-MS hits were predicted using Alpha-Fold2 (37). Protein disorder was predicted using the IUPred3 webserver (38). Both Alpha-Fold2 and IUPred3 predicted disorder in the same regions.

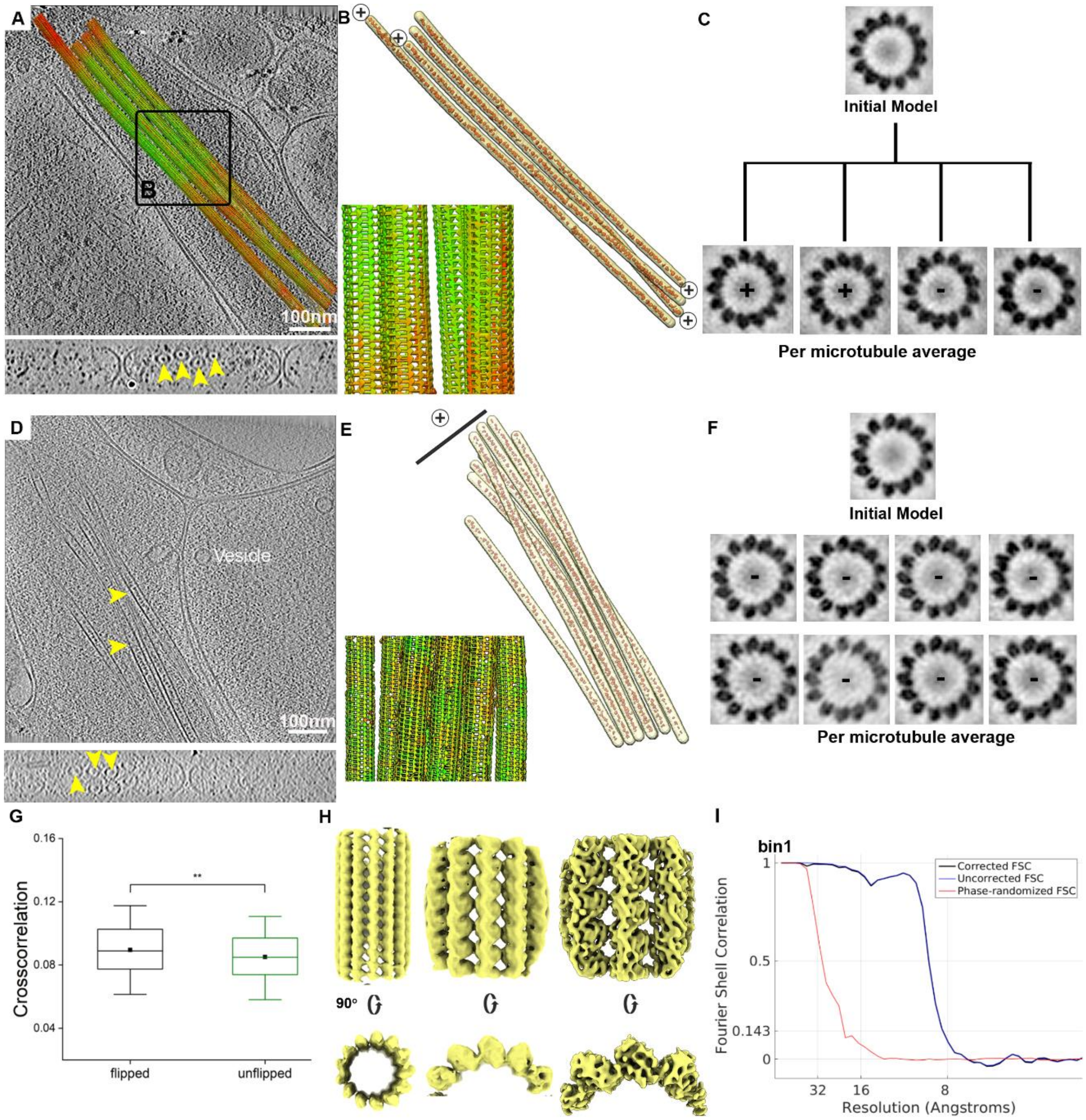


**Fig. S1. Determination of MT polarity and structure exemplified in primary neuron processes**. **A**. top: 9 nm x-y tomographic slice of a neuronal process containing 4 MTs superimposed with lattice map following STA (Methods, also shown in Figure 1A). Green indicate high CCC-values, whereas orange and red indicate lower CCC-values. Bottom: missing wedge compensated, denoised cross-section view shows centrally located lumenal particles (x-z slice; arrowheads). **B**. Segmented MTs with polarity assigned based on the description in C. Bottom left: zoomed in image of the lattice map within the black box in (A) showing different orientations of the subtomograms (represented as arrows). **C**. Individually averaged MTs displaying different polarities indicated by plus/minus sign based on the skew of the tubulin subunits. **D**, **E**, **F**. same as in (A-C**)**

but MTs have uniform polarity orientations. **G.** Comparison of CCC-values between flipped and un-flipped data of the reference used to determine MT polarity, with the mean (black dot). Error bar indicate standard error. Statistical significance **p<0.01, obtained by nonparametric Mann-Whitney test. **H**. Subtomogram averages of the neuronal MT at 4x binning, 2x binning and un-binned resolution (bin 1; pixel size 2.24Å). **I**. Fourier Shell Correlation (FSC) plot of the un-binned subtomogram average obtained using 34906 subtomograms from three tomograms.

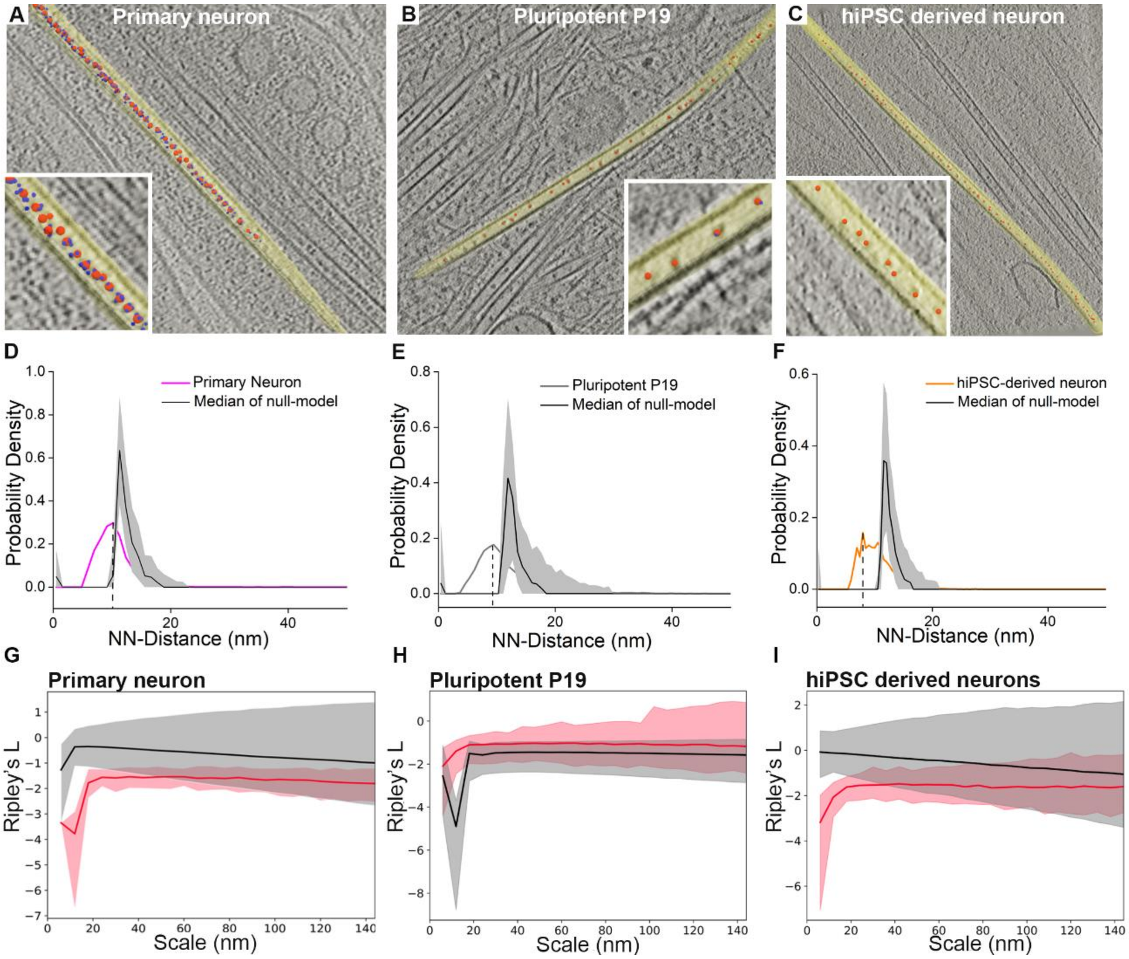


**Fig. S2**. **Template-free detection, picking and organization of lumenal particles. A.** 9 nm thick tomographic slice of a primary neuronal process. Small blue spheres represent local minima obtained after topological simplifications (18), big red spheres are the final picking points representing mean shift clustering centers. Inset shows zoomed in image of the same. **B**. 6.8 nm thick tomographic slice of a pluripotent P19 cellular process. Red dots showing detected particles after topological persistence and mean shift clustering. Inset shows zoomed in image. **C**. Same as in (B) for hiPSC derived neurons, slice thickness 4.25 nm. **D-F.** NN-distance distributions of the lumenal particles in indicated cell types. Null-model simulations (SI Appendix) representing complete spatial randomness are shown in black and shaded gray regions indicate interval of confidence (IC) [5, 95] %. Dashed vertical black lines indicate the most frequent nearest neighbor distances. **G-I**. Univariate 2$^{nd}$ order Ripley's L showing uniform distribution (SI Appendix) of lumenal particles within MTs of the different cell types. Red and black lines indicate medians of the Ripley's L function for the experimental and simulated random distribution, respectively. Shaded areas define the IC [5, 95] %.

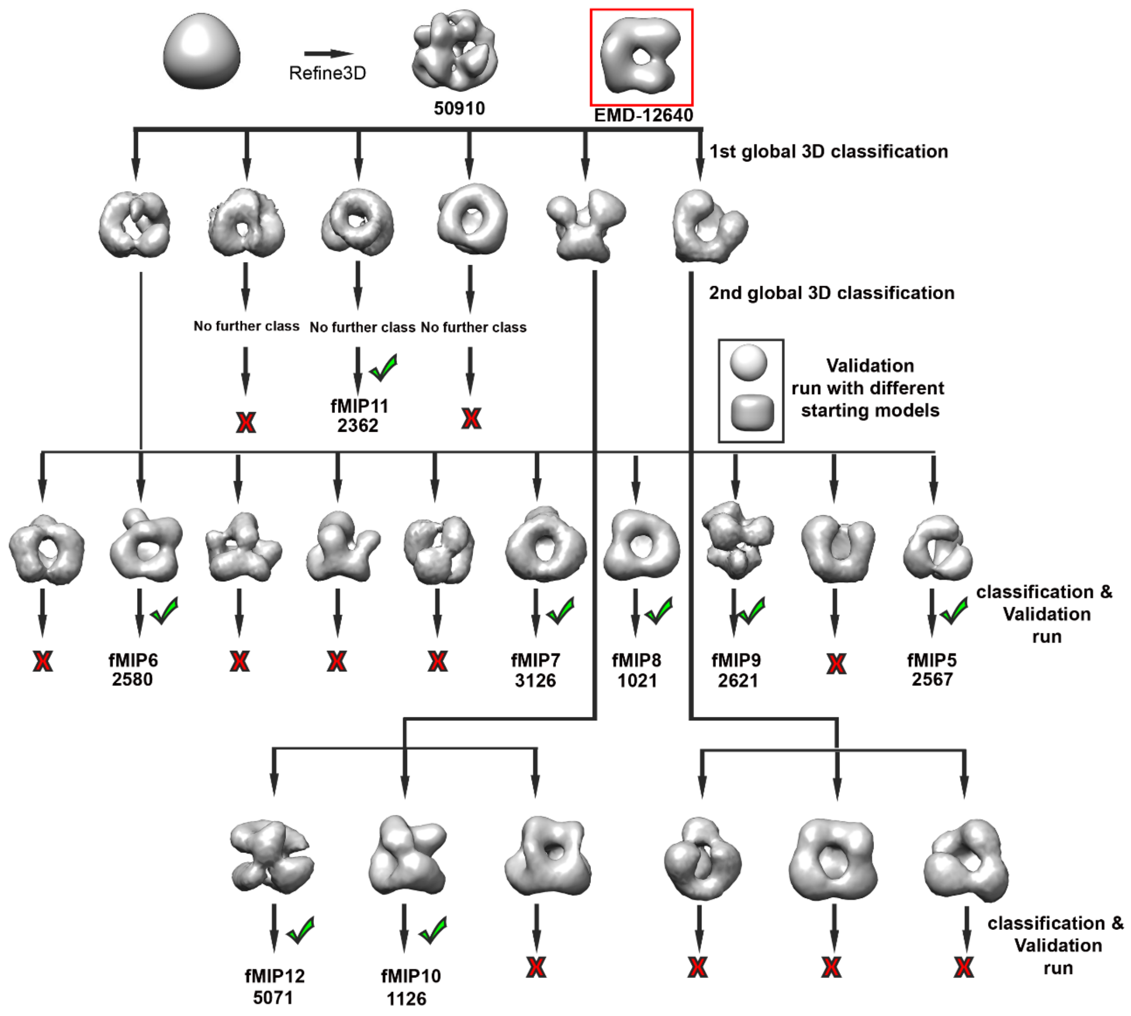


**Fig. S3: Subtomogram averaging and processing workflow of lumenal particles exemplified for primary neuronal cells.** A similar workflow was used for the analysis of MT lumenal particles in all cell types. The initial model (top left) was generated by averaging all picked particles with randomized orientations resulting in sphere-like structure. Initial model was 3D-autorefined in RELION followed by 3D classifications (top middle). Density map obtained by Foster et al. (39) is shown in a red box for comparison, but was not used in the processing. Classification was done sequentially, i.e. outputs of the previous classification were used as input for a new round of classification until no new classes were obtained. The obtained class averages were validated for model bias and optimum-refinement (40) using soft-edged geometric objects (sphere or cylinder) as starting references. For each class average, the corresponding particle list was used for two more separate auto-refinements with sphere and cylindrical starting models. If similar structures are produced in the separate refinements, indicating no model-bias, the class average was accepted (green tick; similarity comparison also exemplified in Fig.S4 B-D). Number of particles belonging to a particular class average is indicated below each accepted average. Number of tomograms, subtomograms and identified classes for each cell type are provided in Table S1.

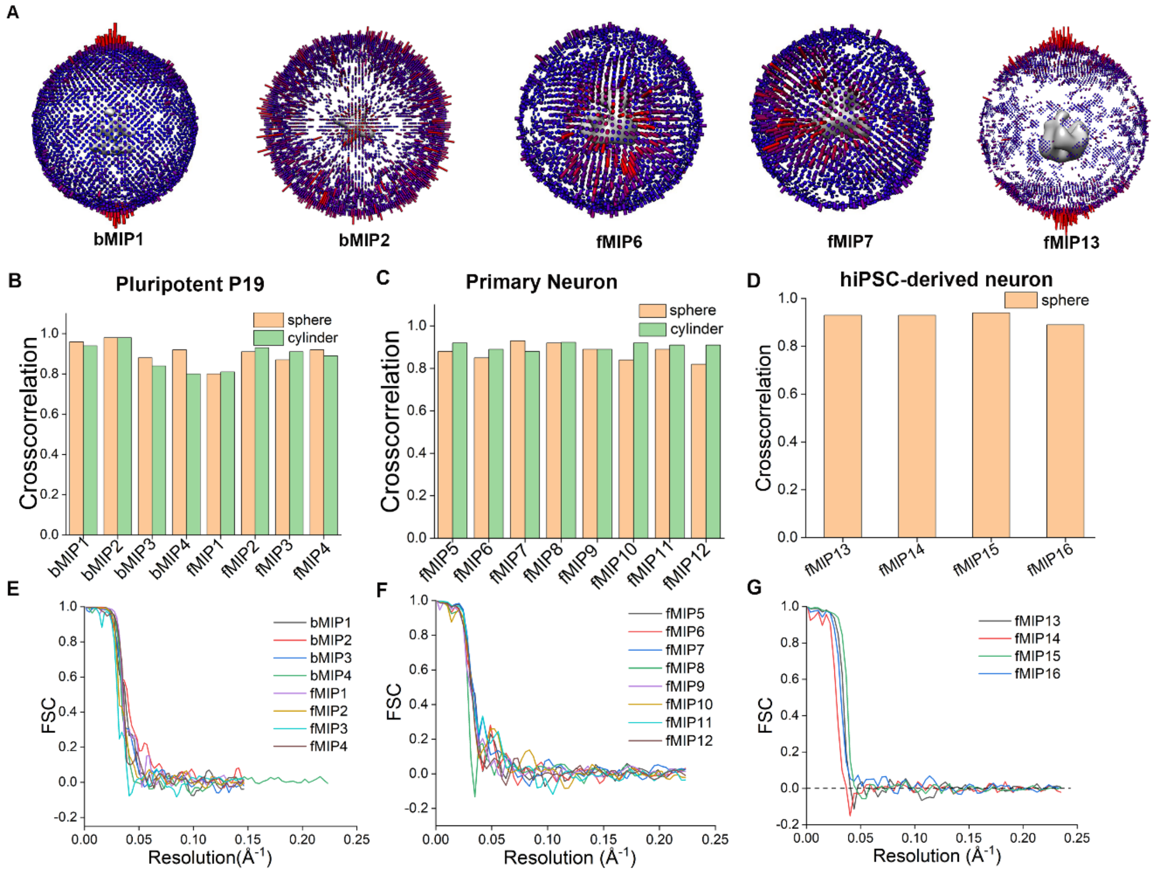


**Fig. S4: Validation of STA-generated lumenal particle maps from various cell lines. A.** Angular distributions obtained from the averages for representative cases. Among all the reported averages, fMIP13 shows modest orientation bias. **B-D.** Similarity between the averages obtained using soft-edged geometric objects (sphere or cylinder) as starting references for refinement and the corresponding averages obtained from RELION starting model, evaluated using cross-correlation coefficient in Chimera for **B**) pluripotent P19 cells, **C**) primary neuron and **D**) hiPSC-derived neurons,. **E-G.** FSC curves of the post-processed averages from **E**) pluripotent P19 cells, **F**) primary neuron and **G**) hiPSC-derived neuronal samples.

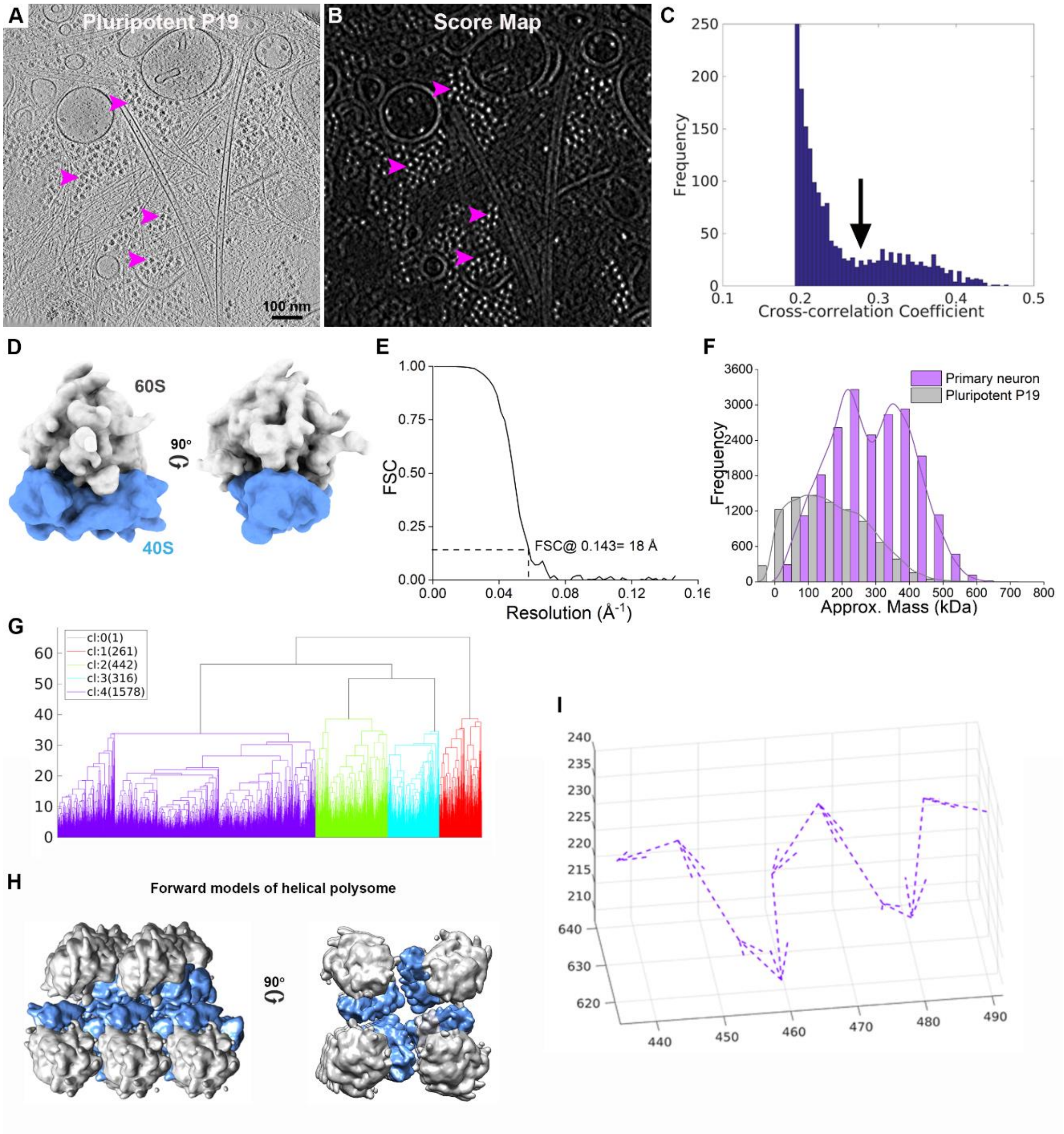


**Fig. S5: Template matching, STA and organization of cellular ribosomes. A.** 6.8 nm tomographic slice of a pluripotent P19 cytoplasm. Examples of ribosomes are indicated by violet arrowheads. **B**. Cross-correlation score map of the same tomogram following template matching in PyTom. Bright spots indicate high scores. Same ribosome clusters as in A are indicated by violet arrowheads. **C**. Histogram of cross-correlation coefficients obtained after template matching. Bimodal distribution indicates false positives (low CCC) and real hits. Black arrow indicates cut-off value chosen to eliminate false positives. **D**. Density map of 80S murine ribosome obtained from 4723 particles by STA in RELION. Large (60S, grey) and small (40S, blue) subunits are indicated. **E**. FSC curve of the 80S ribosome average obtained after RELION post-processing. **F**. Histogram showing approximate mass distributions of the lumenal particles of the indicated samples calculated from the corresponding tomograms. Particle mass was measured using isosurface threshold that corresponds to the molecular weight of the 80S ribosome particles present in the same tomograms. Lines indicate fit to the data distribution. **G**. Hierarchical clustering of transformations of distance matrices calculated from rotations and translations between ribosome pairs. **H**. Representative helical polysome class corresponding to largest cluster (purple) in (G). 60S subunits (gray) face the cytosol. **I**. Vector diagram shows longest helical polysome traced. Arrows indicate translations from n to n+1 ribosome. Axis unit is in pixels, 1 pixel =3.42 Å.

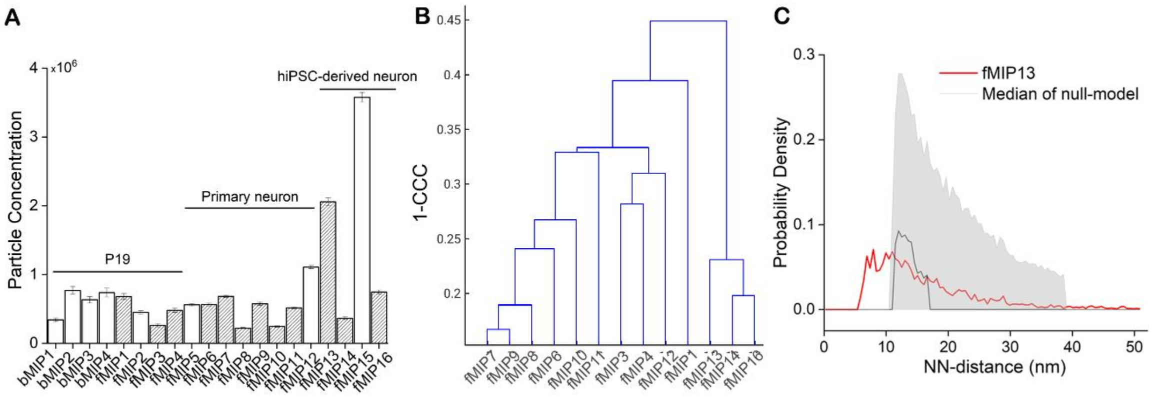


**Fig. S6**: **Lumenal particle abundances and their structural similarity**. **A**. Concentrations of the various class averages obtained from pluripotent P19 cells, primary neuron and hiPSC-derived neuron as indicated. Error bar indicates SEM. Gray bars correspond to the fMIPs shared between all analyzed cell-types possessing ring-like scaffold. Number of tomograms, subtomograms and identified classes for each cell type are provided in Table S1. **B**. Hierarchical clustering tree of the ring-containing fMIPs. All ring-like scaffold containing density maps might represent complexes with variable compositions assembled on the ring-scaffold, or different conformations, determination of which is nontrivial at the obtained low resolution. Therefore, all indicated fMIPs (grey bars) were aligned on the ring-scaffold using rigid-body fitting. Pair-wise cross-correlation values were calculated using the Tom package among all the fMIPs with a limited angular search and evaluated using a hierarchical clustering. Cluster tree shows classes of fMIPs branching only at high CCC values indicating structural similarity. **C**. NN-distance distribution of indicated class average in hiPSC-derived neurons. Null-model simulations (Methods) representing complete spatial randomness are shown in black and shaded gray regions indicate interval of confidence (IC) [5, 95] %.

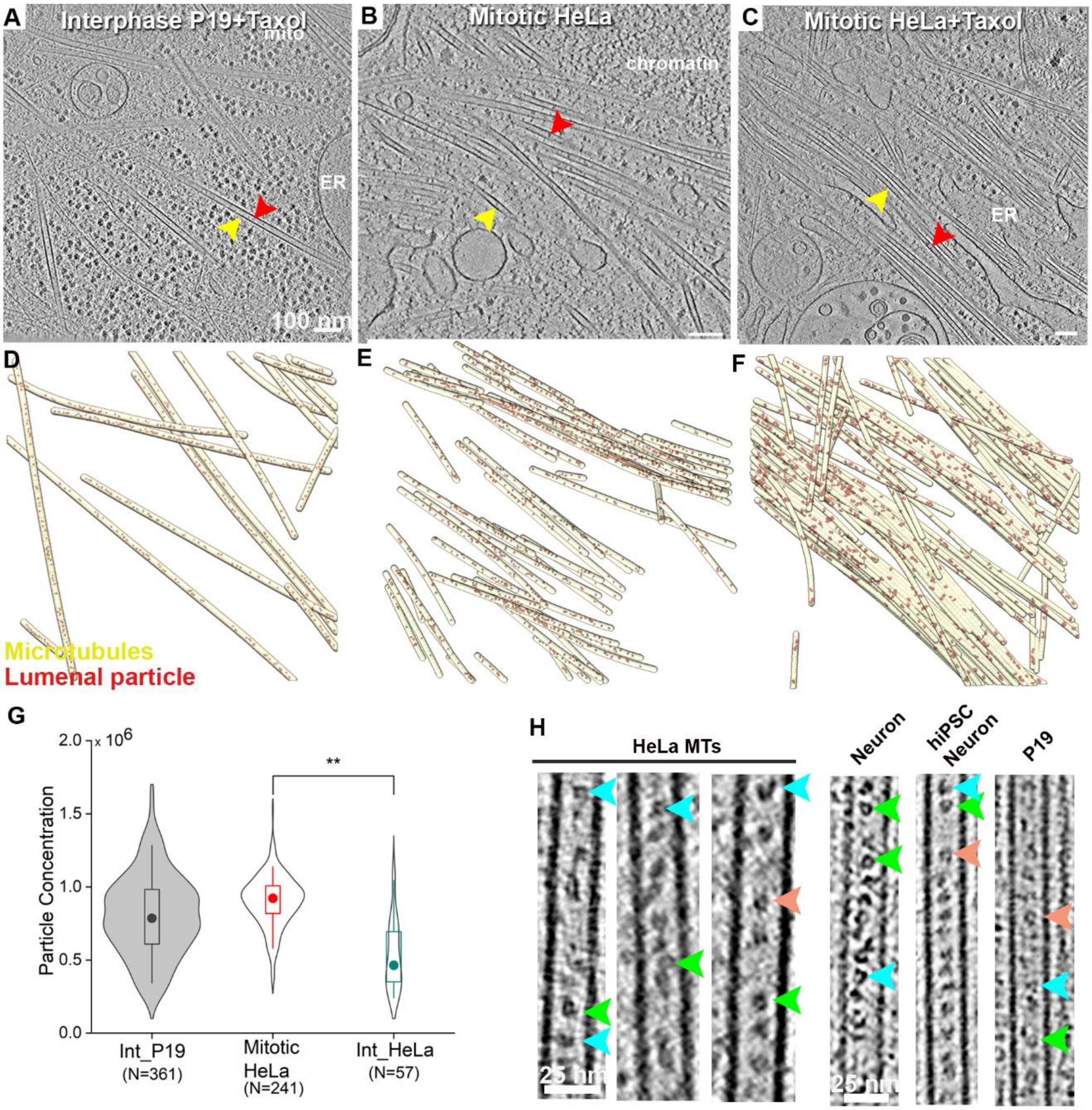


**Fig. S7. Effect of Taxol on lumenal particle abundance in mitotic HeLa cells expressing GFP tagged β–tubulin.** Tomographic slices (top row) and 3D-surface rendered MTs (yellow), along with the lumenal particles (red) of indicated cells (bottom row): **A, D**. Taxol treated pluripotent P19 cell, slice thickness 6.8 nm; **B, E**. mitotic HeLa cell, slice thickness 8.4 nm; **C, F**. Taxol treated mitotic HeLa cell, slice thickness 8.4 nm. **G.** Quantification of the particle concentrations (numbers/μm$^3$ of lumenal volume) in HeLa cells. For comparison, concentrations for P19 and mitotic HeLa from Fig.1I and 3F, respectively are replotted here. Median values are marked by a circle. Asterisks indicate Mann-Whitney test **: significance p<0.01; N, number of MTs. **H**. Morphological similarity between the lumenal particles of all cell types indicated by arrowheads. Same color indicate visually similar particles. Ring-like structure containing lumenal particles are indicated by green arrow head.

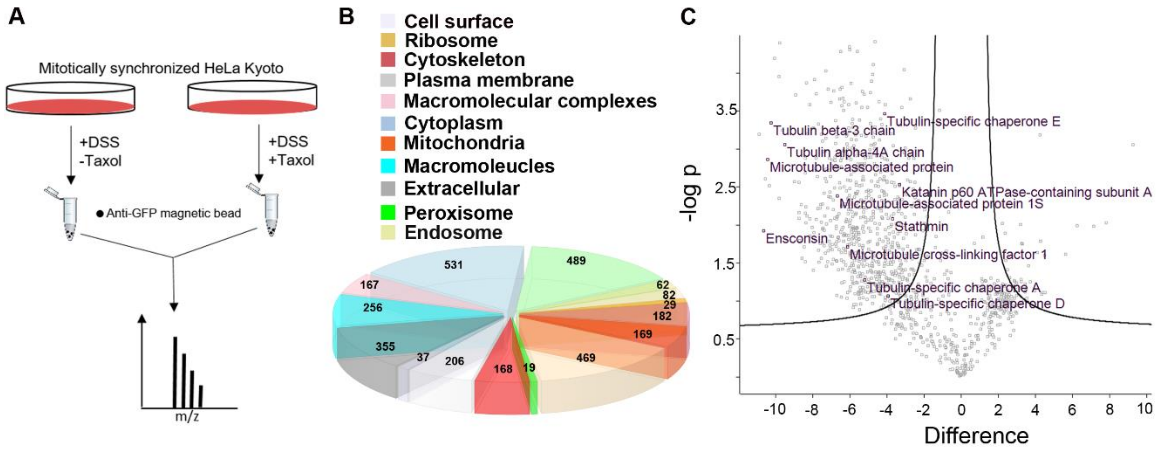


**Fig. S8. AP-MS analysis of MT-interacting proteins in mitotic HeLa cells expressing GFP tagged β–tubulin**. **A**. Experimental strategy for Mass Spectrometry-based identification of lumenal particles from their fold reduction due to Taxol treatment. DSS: Disuccinimidyl suberate. **B**. Pie-chart shows distribution of cellular components based on gene-ontology (GO)-categorization of the affinity purified proteins identified in the Mass spectrometry-based proteomics. Numbers indicate number of protein that belong to each subcategory. 168 proteins are found to be cytoskeleton related. **C**. Volcano plot showing fold change of affinity purified proteins between control and Taxol treated HeLa cell samples. MT-cytoskeleton related top hits indicated by the significant fold ($p<0.05$) change are highlighted. See also **Fig. S9** and **Table S2**.

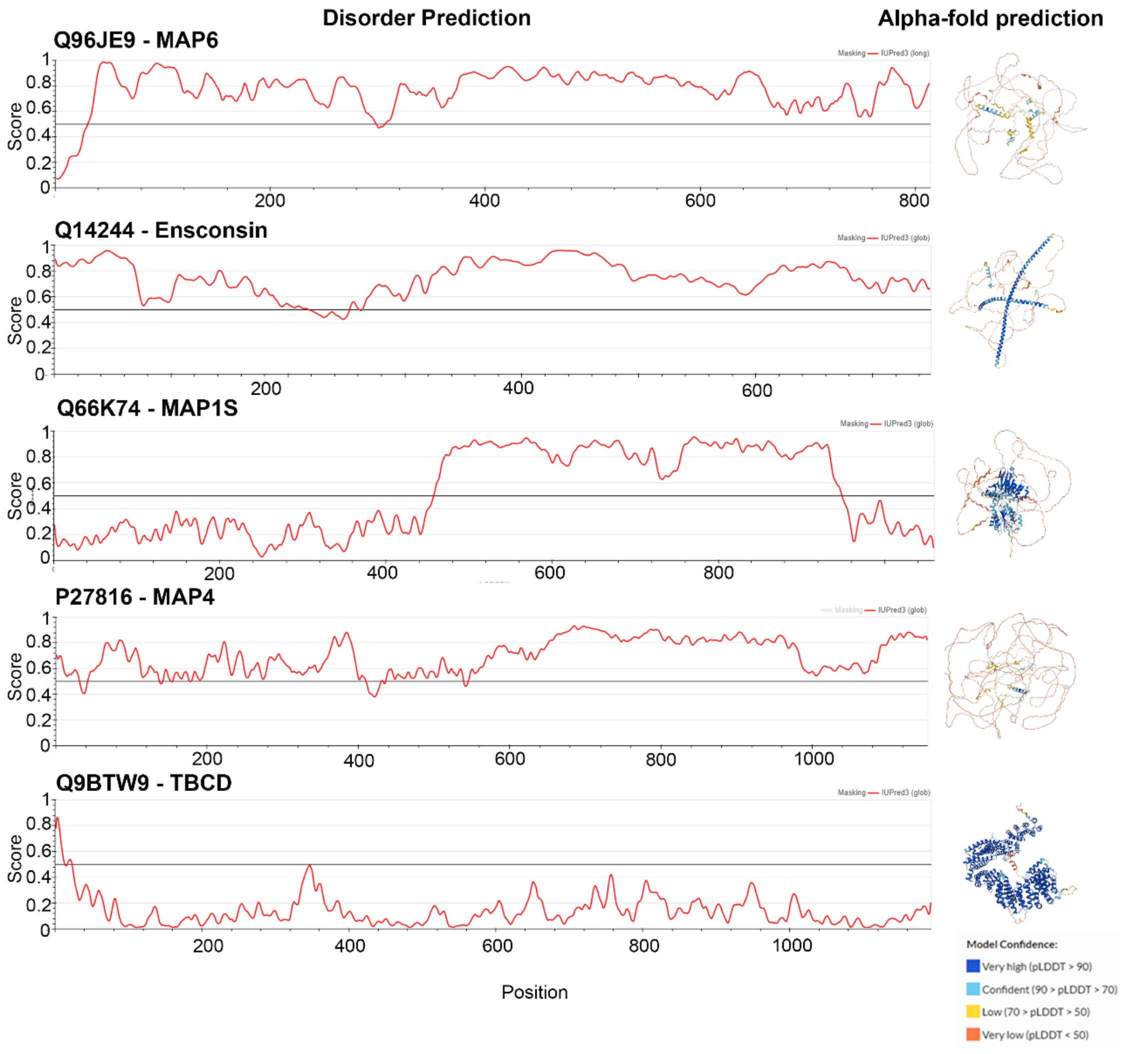


**Fig. S9**: **Protein disorder prediction of select MT-cytoskeleton related hits**. UniProt identifiers are indicated along with the protein IDs (human). Disorder prediction is done with IUPred3 server (38). Red lines indicate prediction score on the scale of 0 to 1. Threshold (black line) is set at default value of 0.5. Scores above the threshold indicate disordered regions. Right panel shows AlphaFold2 (37) predicted structures of the proteins. Color coding indicate per-residue confidence score (pLDDT) in a scale between 0 and 100. All detected MAPs are predicted to be largely unstructured, whereas TBCD has clearly a globular structure. For more comprehensive coverage, see **Table S2**. Although we do not find MAP6, a suggested component of lumenal particles in our AP-MS hits, it is included here for comparison.

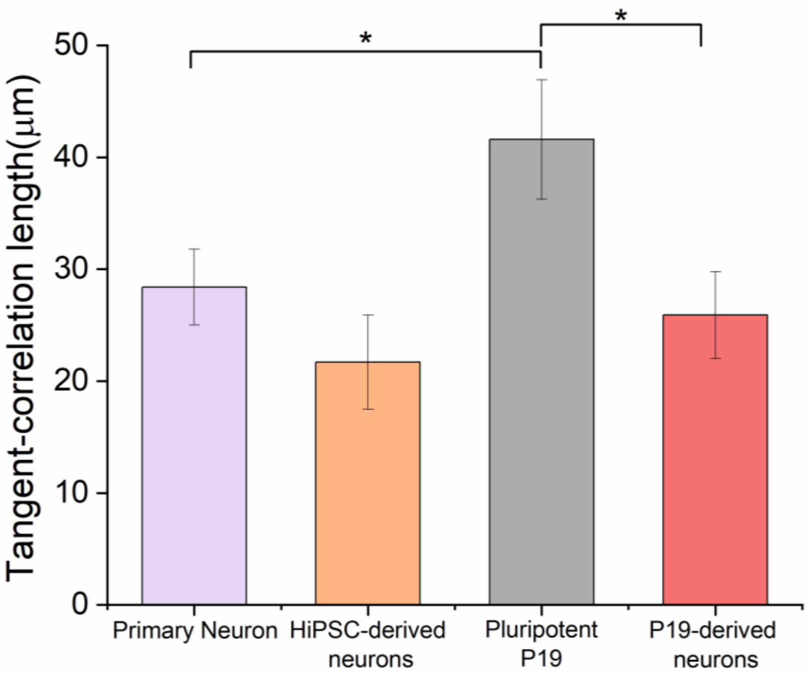


**Fig. S10**: **Quantification of MT curvatures**. Bar graph representing tangent-correlation lengths ($^{a}L_{p}$) of MTs in primary neurons (N=40), hiPSC-derived neurons (N=4), pluripotent P19 cells (N=24) and P19-derived neurons (N=26). N, indicate number of tomograms analyzed. Error bar indicates SEM. Statistical significance *$p<0.05$, obtained by two-sample t-test.

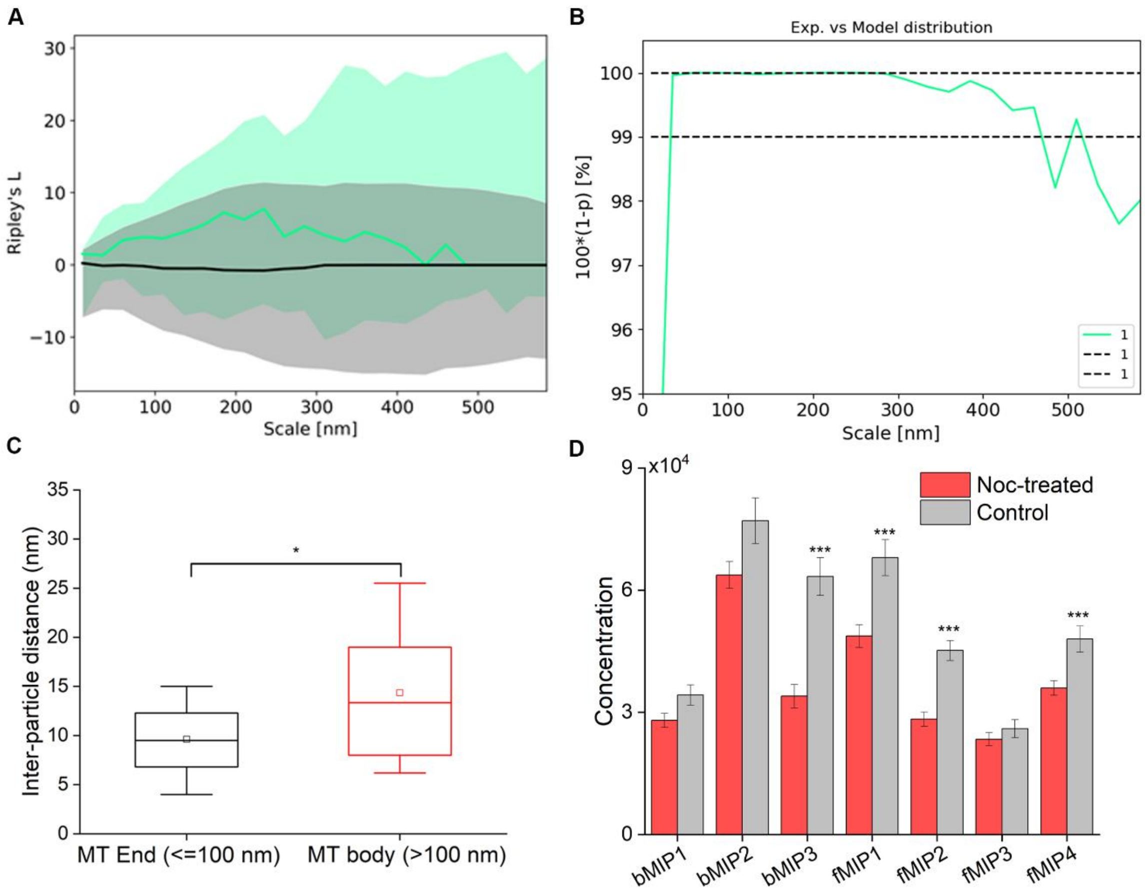


**Fig. S11: Clustering of particles at the MT open ends in Noc-treated pluripotent P19 cells. A.** Experimental (green) vs. random simulated null-model (black). Lines represent the median and shaded areas the IC [5, 95]%. **B**. One-sided Welch test for comparison between the experimental mean (green line) and the simulated one for every scale. Black dashed lines demarcated region indicate area of 99% significance. At a distance >200 nm, significance drops indicating clustering (higher density than random null-model) only occurs at the open end. **C**. Boxplot shows distance between the particles located within 100 nm from the MT open end (black box) and further away (red box). Line indicate median and small rectangle denotes mean value. N=5 MTs, statistical significance by two sample t-test, $p<0.05$. **D**. Concentrations of the indicated class averages in control and Noc-treated pluripotent P19 cells. Error bar indicates SEM. Statistical significance ***$p<0.001$, obtained by two-sample t-test.

**Table S1.** Summary of the data sets from different cell types used for STA.

| | *Murine P19 cells** | *Human iPSC derived neuron* | *Rat primary neuron* | *Global resolution FSC@0.143 (Å)* |
|---|---|---|---|---|
| *#VPP tomograms* | 46 | 8 | - | - |
| *#Defocus tomograms* | 35 | - | 60 | - |
| *#subtomograms* | 44,944 | 8635 | 50910 | - |
| *#bMIP1 subtomograms* | 4411 | - | - | 21.05 |
| *#bMIP2 subtomograms* | 4466 | - | - | 16.69 |
| *#bMIP3 subtomograms* | 3224 | - | - | 21.46 |
| *#bMIP4 subtomograms* | 1519 | - | - | 24.45 |
| *#fMIP1 subtomograms* | 3825 | - | - | 20.12 |
| *#fMIP2 subtomograms* | 2536 | - | - | 23.87 |
| *#fMIP3 subtomograms* | 1653 | - | - | 26.39 |
| *#fMIP4 subtomograms* | 3060 | - | - | 25.91 |
| *#fMIP5 subtomograms* | - | - | 2567 | 25.25 |
| *#fMIP6 subtomograms* | - | - | 2580 | 26.18 |
| *#fMIP7 subtomograms* | - | - | 3126 | 20.92 |
| *#fMIP8 subtomograms* | - | - | 1021 | 32.26 |
| *#fMIP9 subtomograms* | - | - | 2621 | 22.03 |
| *#fMIP10 subtomograms* | - | - | 1126 | 26.81 |
| *#fMIP11 subtomograms* | - | - | 2362 | 28.67 |
| *#fMIP12 subtomograms* | - | - | 5071 | 26.60 |
| *#fMIP13 subtomograms* | - | 2151 | - | 26.46 |
| *#fMIP14 subtomograms* | - | 380 | - | 30.86 |
| *#fMIP15 subtomograms* | - | 3738 | - | 24.51 |
| *#fMIP16 subtomograms* | - | 779 | - | 27.1 |
| *#80S Ribosome subtomograms* | 4723 | - | - | 17.27 |
| *# MT subtomograms* | | | 34906 | 8.19 |

***** In case of pluripotent and differentiated P19 cells, both VPP and defocus data were obtained and processed separately.

**Table S2**. MT cytoskeleton related proteins obtained from the AP-MS experiment with their Alphafold-2 predicted structures. Comments on the structural disorder determined from IUPred3 server also included. Color coding of Alphafold-2 predicted structures indicate per-residue confidence score (pLDDT) in a scale between 0 and 100. Please see Fig.S9 for detail.

| Hits | UniProt ID | Function | Mol. Wt.<br><br>(kDa) | Structural disorder (IUPred3) | Predicted structure (Alphafold-2) |
|---|---|---|---|---|---|
| *Microtubule-associated protein 6 (MAP6) | Q96JE9 | MT stabilization | 86.5 | Highly disordered | 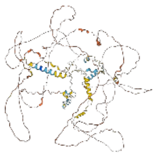 |
| Microtubule-associated protein 1S (MAP1S) | Q66K74 | MT stabilization | 112.2 | Partially disordered | 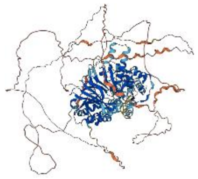 |
| Microtubule-associated protein 4 (MAP4) | P27816 | MT stabilization | 121 | Highly disordered | 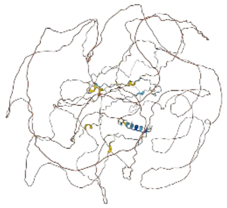 |
| Microtubule-associated protein 7 (MAP7) | Q14244 | MT stabilization | 84 | Highly disordered | 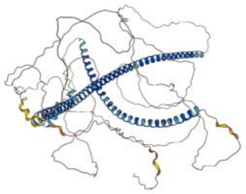 |
| Tubulin Binding Chaperone-E | Q15813 | Tubulin proteostasis | 59.3 | Structured | 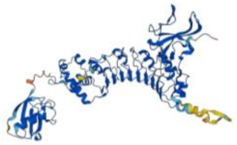 |
| Stathmin | P16949 | MT destabilizatio n | 17.3 | Structured | 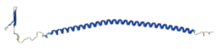 |
| Katanin | O75449 | MT destabilizatio n | 56 | Partially disordered | 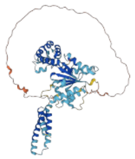 |

| | | | | | |
|---|---|---|---|---|---|
| Kinesin-like protein 3 (KIFC3) | Q9BVG8 | Trafficking | 92 | Structured | 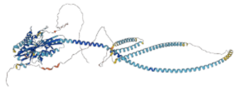 |
| KIF22 | Q14807 | Trafficking | 73.2 | Partially Disordered | 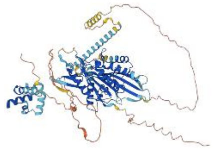 |
| Dynein light chain roadblock-type 1 | Q9NP97 | Cargo recruitment | 10.9 | Structured | 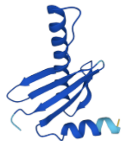 |
| Dynactin subunit 1 | Q14203 | Trafficking | 141.6 | Structured | 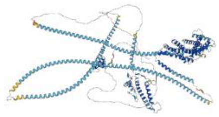 |
| Dynactin subunit2 | Q13561 | Trafficking | 44.2 | Structured | 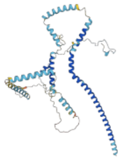 |
| Dynein Light chain 1 | P63167 | Trafficking | 10.3 | Structured | 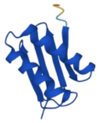 |
| CLIP1 | P30622 | MT polymerizatio n | 162.2 | Partially disordered | 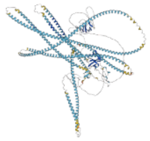 |
| EML4 | Q9HC35 | MT stabilization | 108.9 | Partially disordered | 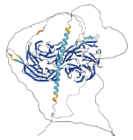 |
| Cytoskeleton-associated protein 2 | Q8WWK9 | MT stabilization | 76.9 | Partially disordered | 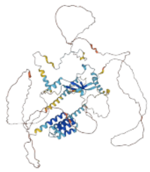 |

| | | | | | |
|---|---|---|---|---|---|
| Cytoskeleton-associated protein 4 | Q9HC35 | ER-MT interaction mediator | 66 | Structured | 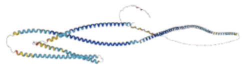 |
| Microtubule cross-linking factor 1 | Q9Y4B5 | MT dynamics regulation | 209 | Partially disordered | 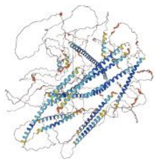 |
| Tubulin beta-3 | Q13509 | MT constituent | 50.4 | Structured | 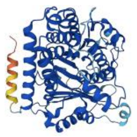 |
| Tubulin Binding Chaperone-A | O75347 | Tubulin proteostasis | 12.8 | Structured | 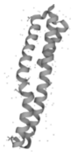 |
| Tubulin Binding Chaperone-D | Q9BTW9 | Tubulin proteostasis | 110 | Structured | 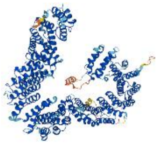 |

*MAP6 is neuron-specific. It is included here only for comparison.

**Supplemental videos titles and legends**

**Movie S1.** Tomographic slices of a primary hippocampal neuronal process showing the ultrastructure of the MT cytoskeleton. Related to Figure 1A, S1A. Color coding, MT: yellow and lumenal particles: red.

**Movie S2.** Tomographic slices of a pluripotent P19 cellular process showing the ultrastructure of the cytoplasm containing MTs. Related to Figure 1B. Color coding, MT: yellow and lumenal particles: red.

**Movie S3.** Tomographic slices of a hiPSC-derived neuronal process showing the ultrastructure of the MT cytoskeleton. Related to Figure 1C. Color coding, MT: yellow and lumenal particles: red.

**Movie S4.** Tomographic slices of a Taxol-treated pluripotent P19 cellular process showing the ultrastructure of the cytoplasm containing MTs. Related to Figure S7A. Color coding, MT: yellow and lumenal particles: red.

**Movie S5.** Tomographic slices of a pluripotent P19 cellular process showing freshly polymerized MTs after Nocodazole treatment and washout. Related to Figure 7G. Color coding, MT: yellow and lumenal particles: red.

### SI References


1. C. Papantoniou, *et al.*, Munc13- and SNAP25-dependent molecular bridges play a key role in synaptic vesicle priming. *Sci. Adv.* **9**, eadf6222 (2023).
2. S. Chakraborty, J. Mahamid, W. Baumeister, Cryoelectron Tomography Reveals Nanoscale Organization of the Cytoskeleton and Its Relation to Microtubule Curvature Inside Cells. *Structure* **28**, 991-1003.e4 (2020).
3. Y. Zhang, *et al.*, Rapid single-step induction of functional neurons from human pluripotent stem cells. *Neuron* **78**, 785–798 (2013).
4. S.-M. Ho, *et al.*, Rapid Ngn2-induction of excitatory neurons from hiPSC-derived neural progenitor cells. *Methods* **101**, 113–124 (2016).
5. M. Toro-Nahuelpan, *et al.*, Tailoring cryo-electron microscopy grids by photo-micropatterning for in-cell structural studies. *Nat. Methods* **17**, 50–54 (2020).
6. J. Mahamid, *et al.*, Visualizing the molecular sociology at the HeLa cell nuclear periphery. *Science* **351**, 969–972 (2016).
7. Y. Fukuda, U. Laugks, V. Lučić, W. Baumeister, R. Danev, Electron cryotomography of vitrified cells with a Volta phase plate. *J. Struct. Biol.* **190**, 143–154 (2015).
8. D. N. Mastronarde, Automated electron microscope tomography using robust prediction of specimen movements. *J. Struct. Biol.* **152**, 36–51 (2005).
9. W. J. H. Hagen, W. Wan, J. A. G. Briggs, Implementation of a cryo-electron tomography tilt-scheme optimized for high resolution subtomogram averaging. *J. Struct. Biol.* **197**, 191–198 (2017).
10. S. Khavnekar, P. Erdmann, W. Wan, TOMOMAN: Streamlining Cryo-electron Tomography and Subtomogram Averaging Workflows Using TOMOgram MANager. *Microsc. Microanal.* **29**, 1020–1020 (2023).
11. S. Q. Zheng, *et al.*, MotionCor2: anisotropic correction of beam-induced motion for improved cryo-electron microscopy. *Nat. Methods* **14**, 331–332 (2017).
12. K. Zhang, Gctf: Real-time CTF determination and correction. *J. Struct. Biol.* **193**, 1–12 (2016).
13. A. Rohou, N. Grigorieff, CTFFIND4: Fast and accurate defocus estimation from electron micrographs. *J. Struct. Biol.* **192**, 216–221 (2015).
14. B. Turoňová, F. K. M. Schur, W. Wan, J. A. G. Briggs, Efficient 3D-CTF correction for cryo-electron tomography using NovaCTF improves subtomogram averaging resolution to 3.4 Å. *J. Struct. Biol.* **199**, 187–195 (2017).
15. D. Tegunov, P. Cramer, Real-time cryo-electron microscopy data preprocessing with Warp. *Nat. Methods* **16**, 1146–1152 (2019).
16. J. R. Kremer, D. N. Mastronarde, J. R. McIntosh, Computer visualization of three-

dimensional image data using IMOD. *J. Struct. Biol.* **116**, 71–76 (1996).

17. Y. T. Liu, *et al.*, Isotropic reconstruction for electron tomography with deep learning. *Nat. Commun.* **13**, 1–17 (2022).
18. A. Martinez-Sanchez, *et al.*, Template-free detection and classification of membrane-bound complexes in cryo-electron tomograms. *Nat. Methods* **17**, 209–216 (2020).
19. D. Comaniciu, P. Meer, Mean shift: A robust approach toward feature space analysis. *IEEE Trans. Pattern Anal. Mach. Intell.* **24**, 603–619 (2002).
20. J. Zivanov, *et al.*, New tools for automated high-resolution cryo-EM structure determination in RELION-3. *Elife* **7:e42166** (2018).
21. E. F. Pettersen, *et al.*, UCSF Chimera?A visualization system for exploratory research and analysis. *J. Comput. Chem.* **25**, 1605–1612 (2004).
22. W. Wan, S. Khavnekar, J. Wagner, P. Erdmann, W. Baumeister, STOPGAP: A Software Package for Subtomogram Averaging and Refinement. *Microsc. Microanal.* **26**, 2516–2516 (2020).
23. K. Qu, *et al.*, Structure and architecture of immature and mature murine leukemia virus capsids. *Proc. Natl. Acad. Sci. U. S. A.* **115**, E11751–E11760 (2018).
24. J. V. Ribeiro, *et al.*, QwikMD — Integrative Molecular Dynamics Toolkit for Novices and Experts. *Sci. Rep.* **6**, 26536 (2016).
25. S. Nickell, *et al.*, TOM software toolbox: acquisition and analysis for electron tomography. *J. Struct. Biol.* **149**, 227–234 (2005).
26. R. Englmeier, S. Pfeffer, F. Förster, Structure of the Human Mitochondrial Ribosome Studied In Situ by Cryoelectron Tomography. *Structure* **25**, 1574-1581.e2 (2017).
27. H. Khatter, A. G. Myasnikov, S. K. Natchiar, B. P. Klaholz, Structure of the human 80S ribosome. *Nature* **520**, 640–645 (2015).
28. T. Hrabe, *et al.*, PyTom: A python-based toolbox for localization of macromolecules in cryo-electron tomograms and subtomogram analysis. *J. Struct. Biol.* **178**, 177–188 (2012).
29. W. Jiang, *et al.*, A transformation clustering algorithm and its application in polyribosomes structural profiling. *Nucleic Acids Res.* **50**, 9001–9011 (2022).
30. A. Martinez-Sanchez, W. Baumeister, V. Lučić, Statistical spatial analysis for cryo-electron tomography. *Comput. Methods Programs Biomed.* **218**, 106693 (2022).
31. E. F. Pettersen, *et al.*, UCSF ChimeraX: Structure visualization for researchers, educators, and developers. *Protein Sci.* **30**, 70–82 (2021).
32. A. Cook, *et al.*, The structure of the nuclear export receptor Cse1 in its cytosolic state reveals a closed conformation incompatible with cargo binding. *Mol. Cell* **18**, 355–367 (2005).
33. S. Nithianantham, *et al.*, Tubulin cofactors and Arl2 are cage-like chaperones that regulate the soluble αβ-tubulin pool for microtubule dynamics. *Elife* **4:e08811** (2015).

34. I. Poser, *et al.*, BAC TransgeneOmics: A high-throughput method for exploration of protein function in mammals. *Nat. Methods* **5**, 409–415 (2008).
35. J. Cox, M. Mann, MaxQuant enables high peptide identification rates, individualized p.p.b.-range mass accuracies and proteome-wide protein quantification. *Nat. Biotechnol.* **26**, 1367–1372 (2008).
36. S. Tyanova, *et al.*, The Perseus computational platform for comprehensive analysis of (prote)omics data. *Nat. Methods* **13**, 731–740 (2016).
37. J. Jumper, *et al.*, Highly accurate protein structure prediction with AlphaFold. *Nature* **596**, 583–589 (2021).
38. G. Erdos, M. Pajkos, Z. Dosztányi, IUPred3: Prediction of protein disorder enhanced with unambiguous experimental annotation and visualization of evolutionary conservation. *Nucleic Acids Res.* **49**, W297–W303 (2021).
39. H. E. Foster, C. Ventura Santos, A. P. Carter, A cryo-ET survey of microtubules and intracellular compartments in mammalian axons. *J. Cell Biol.* **221**, e202103154 (2022).
40. S. H. W. Scheres, Amyloid structure determination in RELION -3.1. *Acta Crystallogr. Sect. D Struct. Biol.* **76**, 94–101 (2020).
